# Massively parallel single molecule tracking of sequence-dependent DNA mismatch repair *in vivo*

**DOI:** 10.1101/2023.01.08.523062

**Authors:** Tunc Kayikcioglu, Jasmin S. Zarb, Sonisilpa Mohapatra, Chang-Ting Lin, James A. London, Kasper D. Hansen, Richard Fishel, Taekjip Ha

## Abstract

Whether due to mutagens or replication errors, DNA mismatches arise spontaneously *in vivo*. Unrepaired mismatches are sources of genetic variation and point mutations which can alter cellular phenotype and cause dysfunction, diseases, and cancer. To understand how diverse mismatches in various sequence contexts are recognized and repaired, we developed a high-throughput sequencing-based approach to track single mismatch repair outcomes *in vivo* and determined the mismatch repair efficiencies of 5682 distinct singly mispaired sequences in *E. coli*. We found that CC mismatches are always poorly repaired, whereas local sequence context is a strong determinant of the hypervariable repair efficiency of TT, AG, and CT mismatches. Single molecule FRET analysis of MutS interactions with mismatched DNA showed that well-repaired mismatches have a higher effective rate of sliding clamp formation. The hypervariable repair of TT mismatches can cause selectively enhanced mutability if a failure to repair would result in synonymous codon change or a conservative amino acid change. Sequence-dependent repair efficiency in *E. coli* can explain the patterns of substitution mutations in mismatch repair-deficient tumors, human cells, and *C. elegans*. Comparison to biophysical and biochemical analyses indicate that DNA physics is the primary determinant of repair efficiency by its impact on the mismatch recognition by MutS.

## Introduction

DNA mismatches occur due to external mutagens, spontaneous chemical changes or misincorporation errors by the DNA replication machinery^1^ and need to be corrected to keep the mutation rates low^2^. Because unrepaired mismatches are sources of genetic variation and point mutations which can alter cellular phenotype and cause dysfunction, diseases, and cancer, we need to understand how diverse mismatches in various sequence contexts are recognized and repaired.

In *E. coli*, the mismatch repair (MMR) system starts with mismatch recognition by protein MutS. MutS then exchanges ADP for ATP creating a conformational change needed to form a sliding clamp, and then recruits and loads MutL sliding clamp to the site (Fig. S1)^3^. MutL sliding clamp engages the nickase MutH, creating a nick at a nearby GATC site. A segment of the nicked strand is removed by the combined action of UvrD helicase and an exonuclease. The single stranded gap thus generated is filled by a DNA polymerase and closed by DNA ligase. MutH nicks the transiently unmethylated nascent strand, thereby favoring the retention of the parental sequence.

One can indirectly infer the sequence dependence of *in vivo* MMR efficiency by comparing the mutation accumulation rates in MMR-capable and deficient cells. However, a mutation can arise via two different mismatch intermediates ^4,5^ and its frequency also depends on the sequence-dependence of the mismatch generation process itself. An alternative approach is based on *in vitro* synthesis of DNA containing a mismatch which is then transformed into a cell. By designing the mismatched construct so that only one of the two strands codes for a functional reporter gene^6-8^, MMR has been shown to have a different efficiency depending on the mismatch type and its sequence context ^6,8-11^. However, it is impractical to use a reporter assay to generate datasets large enough for a systematic investigation of sequence dependence. Covering all eight mismatch types and sequence contexts out to the next-nearest neighbors requires conducting 4^2^·8·4^2^ = 2,048 separate experiments. In addition, a mismatch-mediated substitution must cause an observable phenotypic change, through translation termination or an inactivating mutation, further limiting the sequence space that can be sampled.

Here, we used DNA barcodes to monitor the fate of mismatch library at the single-molecule level *in vivo*. We generated a barcoded vector library, ligated it to a mispaired DNA library, and transformed the resulting plasmid library into *E. coli* (Fig. 1a). As the same barcode is shared only among the descendants of the same ancestral plasmid molecule, sequencing of the replication products reveals single-molecule level information about whether a repair event took place before replication (Fig. 1b). Thus, plasmid replication is used as the clock against which repair efficiencies of diverse mismatches are directly determined in a massively parallel manner.

**Figure 1.**
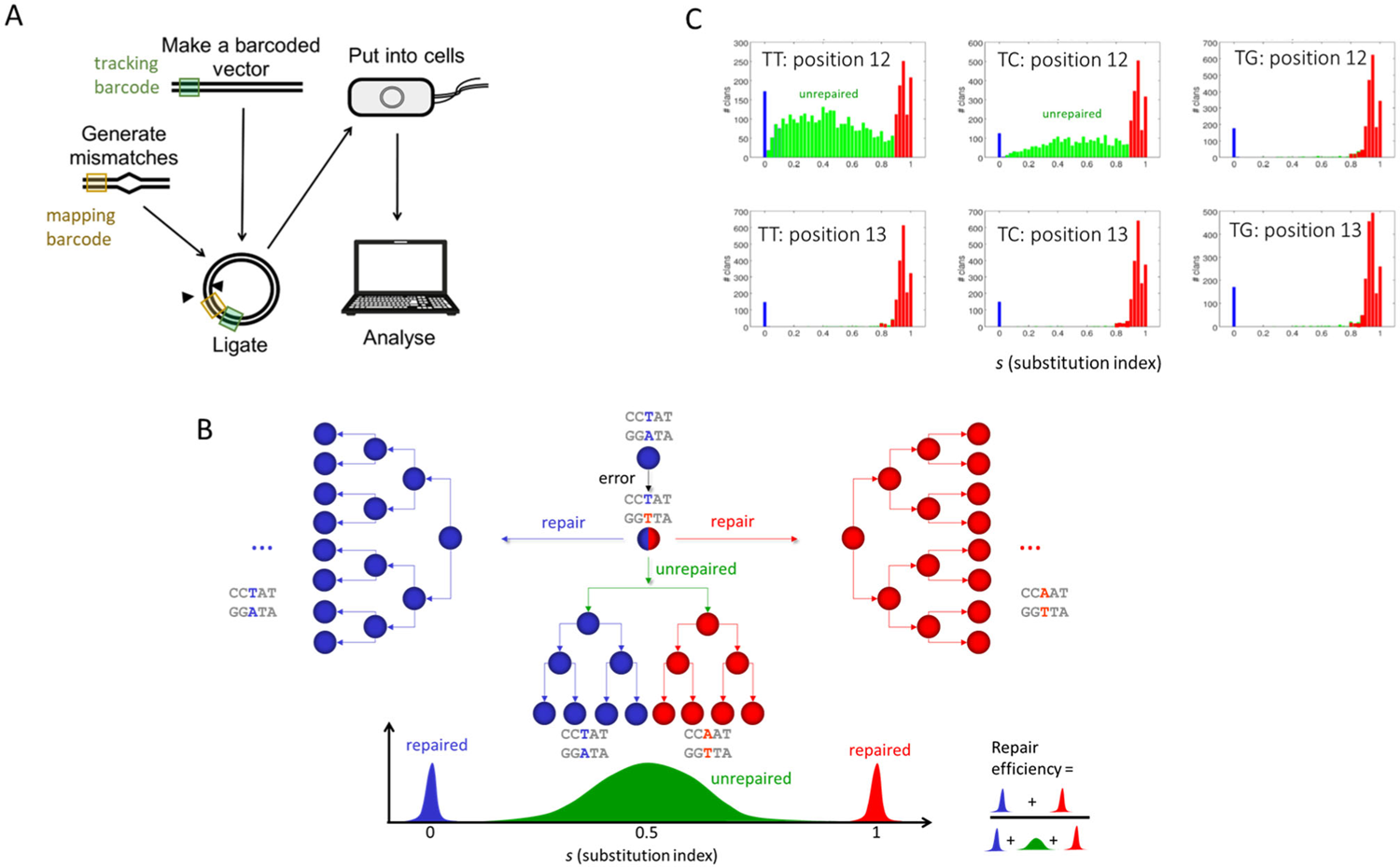
Schematics of MMR-seq and repair efficiency determination. (A) Simplified MMR-seq schematic. A library of mismatch containing DNA containing a mapping barcode, signifying the type and location of a mismatch, is ligated to a plasmid backbone containing the tracking barcode, uniquely identifying the plasmid. The plasmid is put into cells and after rounds of multiplication, the resulting plasmids are harvested and sequenced for analysis. (B) Three outcomes when an error (red) is introduced to create a mismatch (red against blue) (The blue bases and circles represent the initial DNA sequence before replication, while the red bases and circles represent the error that occurs during and after DNA replication resulting in a mismatch). If repair occurs using blue as the template, all of the descendants (or clan members) identified through the unique tracking barcode shared among them, will be ‘blue’ or ‘CCTAA’, and there will be ∼ zero substitution (*s*∼0). If instead repair occurs using red as the template, all of the descendants will be ‘red’ or ‘CCAAA’ with ∼100 % substitution (*s*∼1). If repair does not occur before replication, both blue and red will be found in the same clan, giving an intermediate distribution around *s*∼0.5 (green). (C) Example histograms of *s* for three T-containing mismatches at two neighboring positions. The peaks at low and high s values are colored in blue and red, respectively, and unrepaired clans of intermediate s values are in green.

## Results

### Double barcoding strategy for single molecule tracking of mismatch repair (MMR-seq)

At the single molecule level, individual mismatches should have a binary fate: a particular molecule is either repaired or left unrepaired before the first replication takes place. To track the replication products of mismatch-carrying molecules, we generated plasmids, based on the pUC19 plasmid^12^, each with a unique molecular identifier composed of a randomly generated, 25 bp long ‘tracking barcode’ (Fig. 1a). All plasmids sharing the same tracking barcode can be traced back to a single ancestral plasmid and sequencing reads originating from one of such sister plasmids comprise a ‘clan’.

To examine MMR in all nearest-neighbor sequence contexts, or the ‘trimer’ contexts, while minimizing the total DNA length, we generated a 66 nt long linear De Bruijn sequence, which contains all possible trinucleotides once and only once each. We constructed a DNA library, each member of which deviates from this consensus sequence at exactly one position and amplified this library using a 5’ phosphorylated primer and another primer with a biotin followed by five consecutive phosphothioate bonds at the 5’ end to form an exonuclease-resistant cap (Fig. S2). After selective digestion of the 5’ phosphorylated strand using lambda exonuclease ^13-15^, the surviving ‘variable’ strand is complementary to the ‘constant’ consensus sequence except at one position. In addition, the variable strand contains a ‘mapping barcode’, 6 to 8 bp in length, for mismatch type and position identification. A mismatched library is prepared by extension of a primer containing the constant sequence over the mapping barcode. Finally, the mismatched library is ligated to a PCR-prepared plasmid backbone that also contains the tracking barcode. We transformed this plasmid library into electrocompetent K-12 *E. coli* (BW25113), incubated the transformant bacteria overnight and sequenced at high depth the plasmids extracted. We performed a density-based clustering ^16^ based on the tracking barcodes to organize the sequencing reads into clans that descended from the same ancestral plasmids.

### Repair efficiency determination

As a readout of repair outcome, we determined the ‘substitution index’, *s*, which is the fraction of reads in the same clan that carries the variable strand sequence (Fig. 1b). If the mismatch remains unrepaired until replication, one daughter plasmid inherits the constant strand sequence whereas the other receives the variable strand sequence. Subsequent rounds of plasmid replication would give rise to a heterogeneous clan with an *s* value around 0.5 (Fig. 1b, green, U-type clan, <u>u</u>nrepaired). In contrast, a plasmid that is repaired by keeping the constant strand sequence will carry no variable strand contribution, i.e. *s*=0 (C-type clan, blue, repaired using <u>c</u>onstant strand). Similarly, for a repair that preserves the variable strand, *s*=1 (V-type clan, red, repaired using <u>v</u>ariable strand). However, we found evidence that MMR can initiate but not complete before replication, giving rise to *s* values close to 1 but not 1 (Supplementary Discussion). For our analyses, such outcomes are classified as V-type.

It is instructive to examine a few example histograms of *s* values. For a TT mismatch at position 12 (out of 66 positions of the trimer library), we observe C-type and V-type peaks at 0 and near 1, respectively, plus a broad distribution of intermediate *s* values due to unrepaired U-type clans (Fig. 1c). For a TC mismatch at the same position, we see a lower fraction of U-type, indicating a higher repair efficiency. In contrast, a TG mismatch at the same position did not show any U-type outcome, indicating nearly 100% repair efficiency. At position 13, all three mismatches (TT, TC and TG) are extremely well repaired, with very few U-type events. Therefore, TT and TC mismatches can have very different repair efficiencies depending on the sequence context.

We determined the repair efficiency (η) as the relative population of C- and V-type clans combined. Fig. 2a shows η values in a heatmap grid, constant strand sequence along the horizontal axis and the mismatched base on the variable strand along the vertical axis. Generally, most mismatches are well repaired, with η values averaged over all cases to ∼0.89, with notable exceptions: for example, CC mismatches are always poorly repaired, and other mismatches TT, TC and AG mismatches showed hypervariability in repair efficiency.

**Figure 2.**
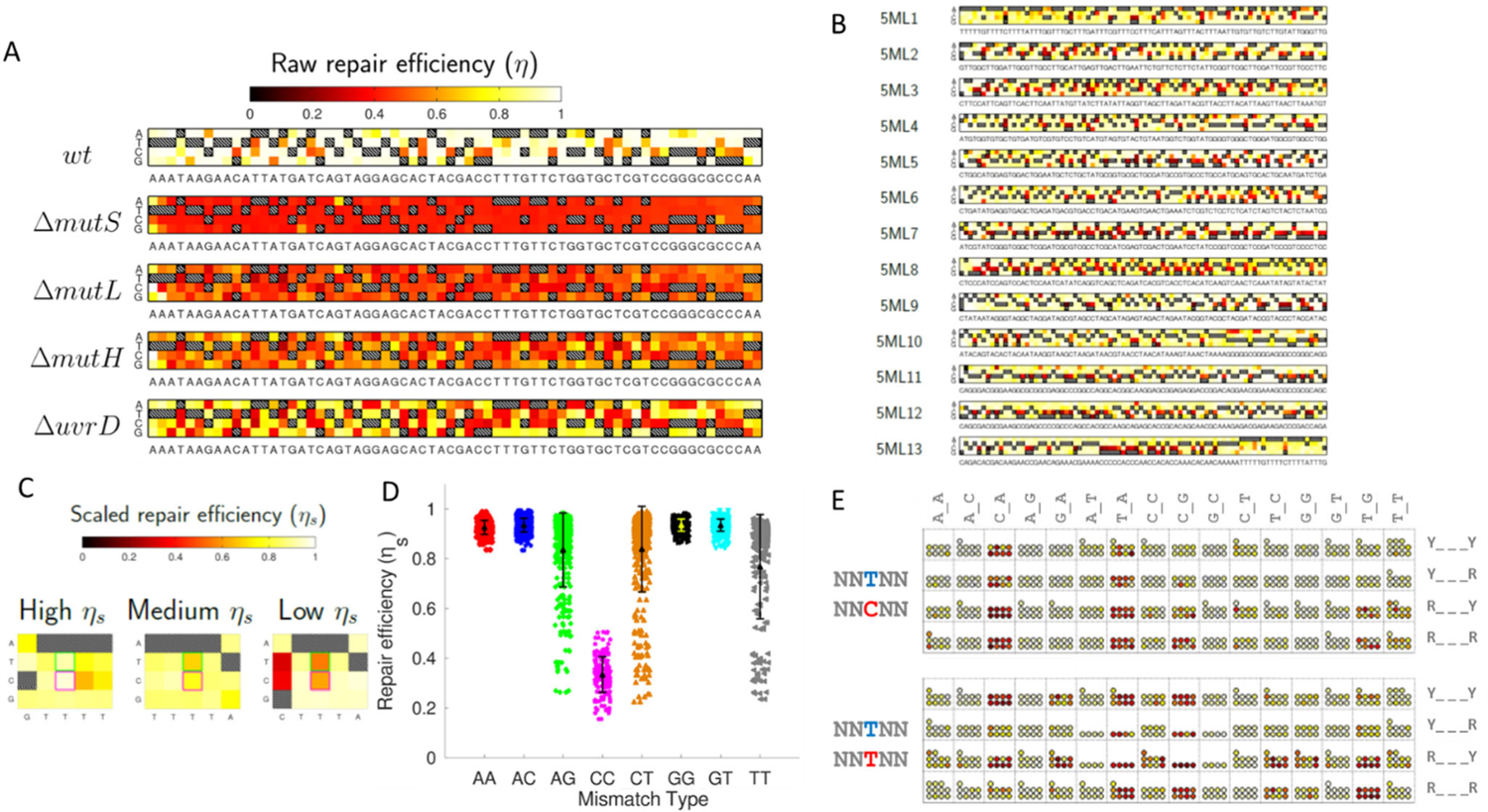
MMR efficiencies for trimer and pentamer libraries. (A) Heat maps of raw repair efficiency values for De Bruijn trimer library from wild type (wt) and deletion strains for *mutS, mutL, mutH* and *uvrD*. The horizontal axis shows the nucleotides on the constant strand and the vertical axis shows nucleotides on the variable strand. Dashed squares are for intact base pairs. (B) Heat maps for De Brujin pentamer library divided into 13 sub-libraries, 5ML1-5ML13. (C) Three pentameric contexts with very different repair efficiencies for TT and GT mismatches even when they share the same trimeric context. (D) Scaled repair efficiency for eight types of mismatches. Each datapoint is for a single pentamer context. Mean and standard deviation are shown in a symbol and error bar. (E) Heat map of scaled repair efficiency for TC mismatch (top) and TT mismatch (bottom), grouped according to the nearest surrounding nucleotides (columns) and the next nearest nucleotides (row). Y for pyrimidines (C or T) and R for purines (A or G).

To confirm that η values reflect the efficiencies of mismatch repair, we performed MMR-seq of the same trimer library but using deletion strains for MutS, MutL, MutH or UvrD. All mutant strains showed greatly reduced η values (Fig. 2a), validating our MMR-seq approach. Interestingly, the averaged η values gradually recovered as more downstream genes are deleted (mean ± SE = 0.89 ± 0.01 for wt, 0.43 ± 0.00 for *mutS*, 0.49 ± 0.00 for *mutL*, 0.54 ± 0.01 for *mutH* and 0.64 ± 0.01 for *uvrD*). Because MutS is essential for mismatch repair, the apparent nonzero η value for *mutS* must be due to some daughter plasmids not producing offspring, giving the appearance of repair. Therefore, we rescaled the repair efficiency so that the scaled repair efficiency, η_s_, is ≈0 for *mutS*.

Why does the apparent repair efficiency recover as more downstream MMR genes are deleted? One possibility is that there are parallel MMR pathways, for example, requiring *mutS* and *mutL* but not *mutH* or *uvrD*. We prefer another explanation where MutS, after detecting the mismatch and launching itself onto DNA, may interfere with replication of one of the two daughter strands either by itself or through the action of other MMR proteins, possibly nicking by MutH, thereby increasing the apparent repair efficiency, which is measured relative to replication.

### Hypervariable repair of AG, CT, TT mismatches

Next, we designed a pentamer library that includes single mismatches in all combinations of nearest neighborhood and second nearest neighborhood, originating from the 1028 bp long, pentamer De Bruijn sequence. Because of length limitations in DNA library synthesis, this pentamer library was divided into 13 sub-libraries of 84 nt length each and MMR-seq was performed on each sub-library (Fig. 2b). Fig. 2c shows an example of three sequence contexts that are identical up to the nearest neighbors (GTTTT, TTTTA, CTTTA). A visual inspection of three CT mismatches (pink box) and three TT mismatches (green box) reveals a striking variation in repair efficiency, showing that the repair is influenced by a local sequence context that goes beyond the nearest neighbors.

Accounting for the symmetry between the two DNA strands, there are 8 possible mismatches (AA, AC, AG, CC, CT, GG, GT and TT). Essentially all CC mismatches were repaired with a low efficiency (η_s_ = 0.35±0.13, s.d.), whereas virtually all AA, AC, GG, and GT mismatches were efficiently repaired irrespective of the sequence context (η_s_ = 0.92±0.07, 0.93±0.04, 0.94±0.04, 0.93±0.04, respectively) (Fig. 2d). In contrast, we observed a wide range of repair efficiencies for AG, CT and TT mismatches (η_s_ = 0.81±0.17, 0.82±0.19, 0.75±0.22, respectively). Using a heptamer library that samples a limited set of sequence context, we also found evidence that sequences beyond the second nearest neighbor can, in some cases, influence the repair efficiency (Fig. S3).

We examined in depth the sequence context dependence of hypervariable repair efficiencies of CT, TT and AG mismatches by tabulating the results according to the identities of neighboring nucleotides (Fig. 2e and Fig. S4). By representing each occurrence of a mismatch by a bead, colored according to η_s_, we can discern patterns. For example, a TC mismatch is almost always poorly repaired (dark red beads) if T is preceded by a pyrimidine and is followed by A regardless of the second nearest neighbors (Fig. 3e, top). In some cases, the next nearest neighbors also determine the outcome in a systematic manner. For example, a TT mismatch in the GTA context is well repaired only when GTA is followed by a purine but not pyrimidine (Fig. 3e, bottom). Overall, the emerging patterns suggest that the identities of the nearest and next neighbors surrounding a mismatch can be used to predict the repair efficiency reasonably well, despite some exceptions we noted earlier.

**Figure 3.**
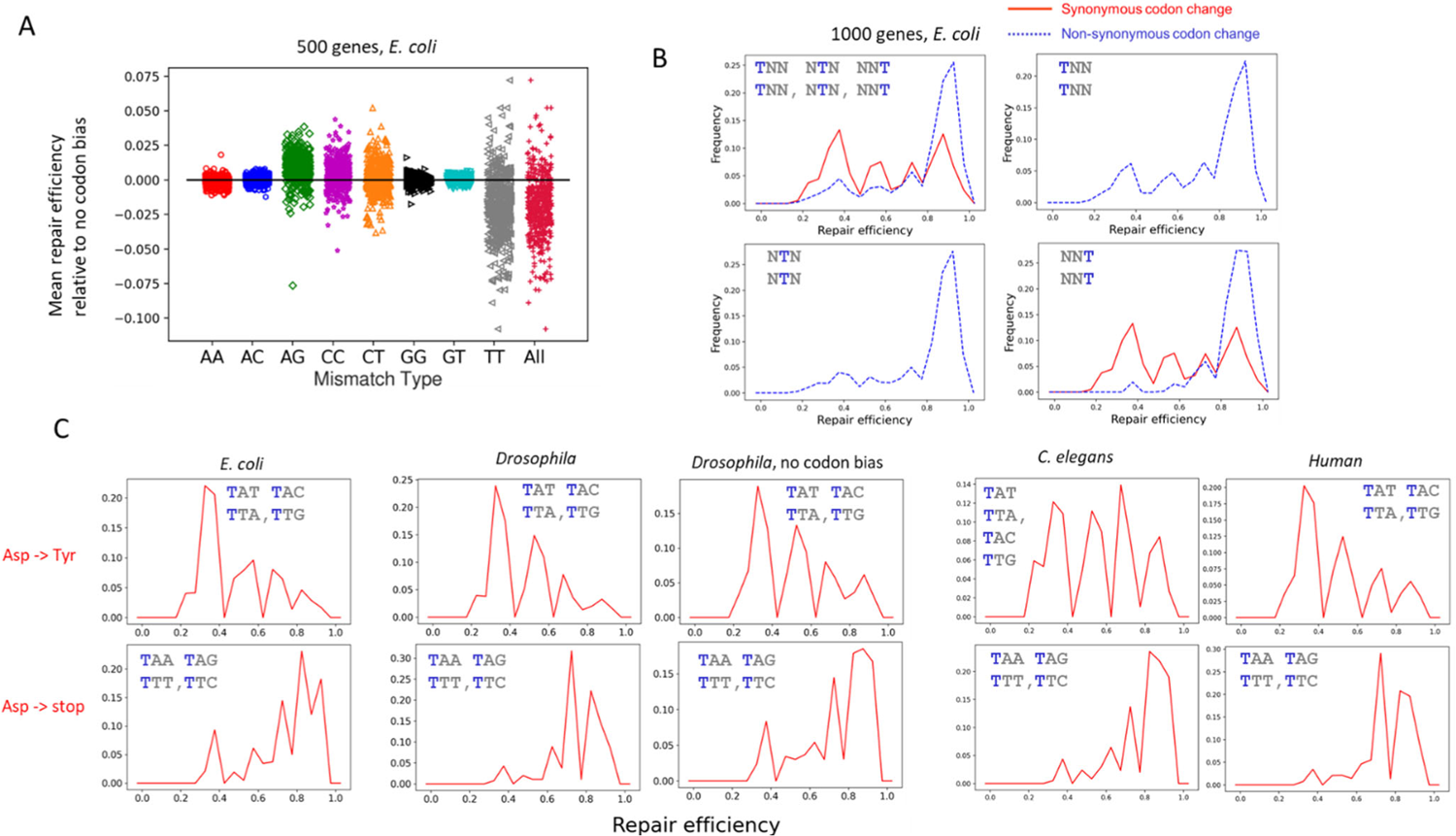
Context-dependent repair and selectively enhanced mutability. (A) Repair efficiency averaged over all possible mismatches of a given type in an *E. coli gene* is predicted relative to the same gene but removing the codon bias. Contributions from TT mismatches dominate the overall behavior (labeled “All” in the plot) where the existing sequence context is more mutable compared to the case without any codon bias. Analysis was done for 500 *E. coli* genes. (B) Repair efficiency histograms for TT mismatches grouped into the cases where a failure to repair would cause a non-synonymous codon change (blue dashed) and other cases where a failure to repair would cause a synonymous codon change (red). In addition, histograms for TT mismatches in positions 1, 2 or 3 in a codon are separately plotted. 1000 *E. coli* genes were included. (C) Repair efficiency histograms for TT mismatches in position 1 of a codon for Asp in sequence contexts where a failure to repair would cause a conservative amino acid change (Asp > Tyr, top) or nonsense codon change (Asp > stop, bottom). In both *E. coli* and *Drosophila*, mismatches that would lead to a generation of a stop codon are repaired more efficiently. In *Drosophila*, randomizing codons without codon bias reduces the contrast.

We also performed MMR-seq on a native DNA sequence from *E. coli*, 490 bp of the *rpoH* gene, including a 5’ regulatory segment, promoter, translation start site, and sequence coding for the N-terminal 84 residues of σ_32_. Again, AG, CT and TT mismatches gave the largest variations (sd = 0.18, 0.21, 0.25, respectively) (Fig. S5), showing their hypervariability in repair efficiencies is not specific to the non-biological De Bruijn sequences.

A previous surface plasmon resonance (SPR) study examined MutS binding to a set of mismatches ^17^. We plotted the predicted η_s_ values of the sequences used in their study versus the dissociation constant *K*_d_ values determined using SPR (Fig. S6a). For all difficult-to-repair mismatches with low η_s_ values, MutS has high *K*_d_ values, showing indeed that lower MutS binding affinity is a common property shared among mismatches that are difficult to repair. Comparing with the biochemical data on human MutS homologs, we found that sequences with low η_s_ values have low *k*_cat_ values for ATP hydrolysis ^18^ (Fig. S6b). Both comparisons indicate that sequence dependent MMR repair efficiency *in vivo* primarily originates from MutS-mismatch interactions.

We next asked if the context-dependent mismatch repair we characterized here influenced the gene sequences. When we randomized synonymous codon choices for 500 *E. coli* protein coding sequences, thereby eliminating codon bias, we observed an increase in the predicted MMR efficiency averaged over all possible mismatches (Fig. 3a). This effect is primarily driven by TT mismatches (Fig. 3a) and is true for Drosophila protein coding sequences (Fig. S7). Although the sign and magnitude of this effect are organism dependent, in all cases examined, the effect is dominated by TT mismatches (Fig. S7). Therefore, the organism-specific codon bias appears to influence gene mutability, mainly exploiting the hypervariable repair of TT mismatches.

Interestingly, the histogram of predicted repair efficiencies of TT mismatches for all occurrences of thymine in protein coding genes is strongly peaked at a high repair efficiency value if a failure to repair the mismatch would result in a change in the amino acid encoded (Fig. 3b for *E. coli*, Fig. S7 for *Drosophila* and *C. elegans*). In contrast, if the sequence context is such that a failure to repair would cause a synonymous codon change, the histogram greatly shifts to lower repair efficiencies (Fig. 3b), suggesting the context dependence of TT mismatch repair may have evolved to preserve the identities of the amino acids encoded while allowing higher mutability for synonymous codon changes. TT mismatches can cause a T>A mutation, which always results in nonsynonymous codon changes if occurring in position 1 or position 2 of a three-letter codon. As a result, the synonymous vs. non-synonymous contrast in repair efficiency is even more pronounced if we restrict the comparison to only the position 3 where a T>A mutation can also result in synonymous codon change (Fig. 3b, Fig. S7). We speculate the much lower repair efficiencies of TT mismatches when failure to repair does not change amino acid may provide an opportunity to optimize other properties of gene sequences such as mRNA secondary structures, codon usage, and DNA mechanics which may affect gene expression.

Another interesting example of a selectively enhanced mutability is that of TT mismatch in position 1 of codons AAU, AAC, AAA and AAG. Predicted repair efficiencies are much lower for AAT/AAC which, if unrepaired, would cause a more conservative, polar to polar amino acid change (asparagine to tyrosine) than for AAA/AAG which will convert lysine to UAA/UAG stop codons in all four organisms examined (Fig. 3c). Further, removing codon bias reduces the contrast in repair efficiencies (Fig. 3c). We also note that this is also another example where the identity of the second nearest neighbor matters; a pyrimidine, T/C, imparts a much lower repair efficiency than a purine, A/G, when it is two base pairs away from the mismatch.

### Single molecule FRET analysis *in vitro* reproduces sequence dependence of MMR *in vivo*

In *E. coli*, after MutS binds a mismatch, it exchanges ADP to ATP and undergoes a conformational change to form a sliding clamp which then recruits MutL (Fig. S1). We adopted the single molecule FRET-based MutS assay ^19^ for six different mismatched DNA whose repair efficiencies *in vivo* have been determined using MMR-seq, with η_s_ ranging from 0.49 to 0.98. MutS was labeled with Cy3 at the N-terminus using a sortase and DNA was labeled with Cy5 at a site 5-8 bp from the mismatch so that MutS binding to a surface-tethered mismatched DNA is observed as a sudden appearance of fluorescence signal with a high FRET efficiency *E*. (Fig. 4a and 4b). Binding was mismatch-dependent because the frequency of binding ranged from 0.02 to 0.12 binding events/s, depending on the mismatched sequence, in comparison to 0.01 binding events/s for a control DNA without a mismatch. Sliding clamp formation and subsequent sliding on DNA is observed as a reduction in *E*, and if MutS slides off the DNA, fluorescence signal abruptly disappears (Fig. 4a and 4b). The same mismatched DNA is observed to recruit multiple MutS molecules in rapid succession as they slide off the mismatch to make it available for the next MutS (Fig. 4b). We modeled the process using MutS (**S**) binding to a mismatch (**M**) with the rate constant *k*_1_ to form **SM, SM** disassembling at the rate *k*_-1_, **SM** converting into **S**^**\***^**M** where **S**^**\***^ denotes MutS sliding clamp, and finally **S**^**\***^ sliding off the DNA through the free end at the rate *k*_3_ (Fig. 4a). From dwell time analyses, we found that *k*_2_ and *k*_3_ are independent of mismatch sequence whereas *k*_1_ increased and *k*_2_ decreased with increasing η_s_ (Fig. 4c, Fig. S8). The duration of the low FRET state after sliding clamp formation is 0.95 s on average, longer than the 0.01 s that it takes for MutS to slide off the 66 bp DNA for a sliding clamp of diffusion coefficient of 3.72 10^5^ bp^2^/s or 0.043 μm^2^/s ^20^. Therefore, the low FRET state may not be entirely attributable to **S**^**\***^ sliding on the DNA and may also include a static complex of **S**^**\***^**M**. Regardless of what happens after sliding clamps formation, we can determine the effective rate of sliding clamp formation, *k*_clamp_ = *k*_1_\**k*_2_/[*k*_2_+*k*_-1_]. *k*_clamp_ displayed a steep monotonic dependence on *η*_s_, showing that much of MMR efficiency variability *in vivo* can be explained by variations in *k*_1_ and *k*_-1_.

**Figure 4.**
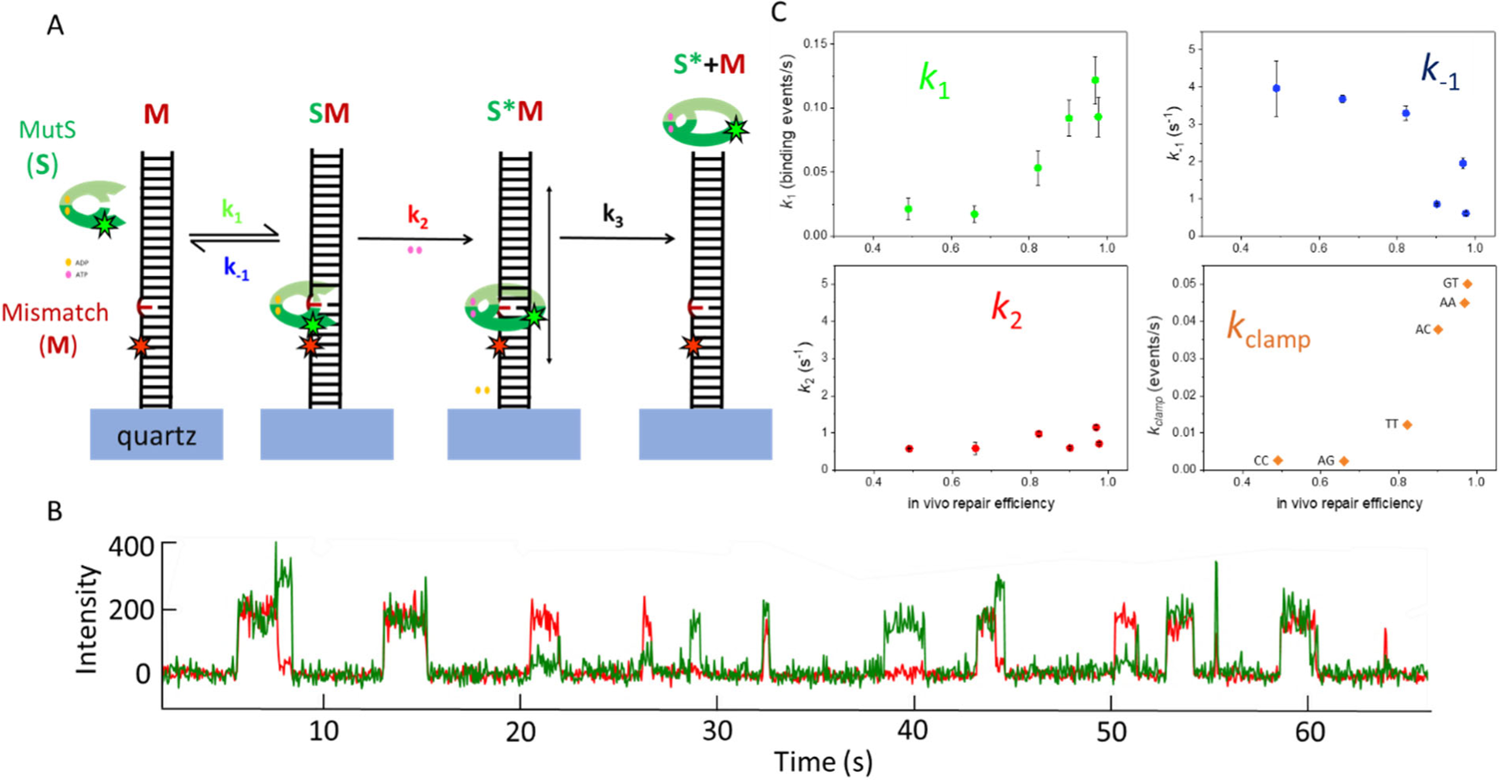
Single molecule analysis of mismatch recognition and sliding clamp formation by MutS. *In vitro* single molecule FRET characterization of MutS’s binding characteristics supports *in vivo* MMR repair trends. (A) Model of MutS’s search, recognition, and sliding clamp formation for a mismatched DNA. *E. coli* MutS N-terminally labeled with a donor (Cy3) initially scans the DNA in search of a mismatched base pair. MutS will bind to the mismatched base pair (highlighted in red) with a rate constant of *k*_1_. MutS can either dissociate from the mismatch base pair with a rate constant of *k*_-1_ or can undergo a conformational change, with a rate constant *k*_2_, by exchanging ADP for ATP, resulting in a sliding clamp. Once a sliding clamp is formed, MutS will slide off the (not end blocked) DNA with a rate constant of *k*_3._ (B) MutS binding to a mismatch and forming a conformational change. A representative trace of MutS’s binding characteristics on a GT mismatched DNA in the presences of 1 mM ATP. Cy3 labeled MutS binds to a mismatched base pair that is located 5-8 bp away from a Cy5 dye, resulting in an initial high FRET state event. MutS then undergoes a pause as it recognizes the mismatch and forms a conformational change, resulting in a change from the high to low FRET state. (C). In the presence of 1 mM ATP, MutS’s *in vitro* binding rate constants, *k*_1_ and *k*_-1_, correlate with the *in vivo* mismatch repair efficiencies reproducing the sequence dependence of *in vivo* mismatch repair characteristics, as shown by the effective rate (*k*_clamp_). However, rate constant, *k*_*2*,_ minimally contributes to MutS’s *in vitro* ability to distinguish among mismatches with different *in vivo* mismatch repair efficiency. Each *k*_clamp_ point is labeled with the corresponding mismatch associated for that data point and repair efficiency.

### Mutational signatures upon MMR inactivation in human cancer and *C. elegans* vs MMR efficiency in *E. coli*

Tumors with defective mismatch repair show characteristic frequencies of single base substitutions that are dependent on the nearest neighbors^21^. The resulting ‘mutational signature’, which is a tabulation of relative frequencies of six types of single base substitution in all possible trimer contexts (6×16 =96 values, accounting for symmetries) ^21^, is likely a reflection of mutations that are normally avoided by a functional MMR system (MMR+) when corresponding mismatches are formed, for example, due to an error by a polymerase. For MMR deficient tumors (MMR-), the frequency of mutations that occur via mismatches that are normally well repaired should increase. Indeed, for mutations that are over-represented in MMR-tumors compared to MMR+ tumors, the corresponding mismatches showed high repair efficiency in *E coli* and vice versa (Fig. 5a). Note that a given point mutation can occur through either of two intermediates, for example C>A mutation can occur through AG or CT mismatch, so we grouped them and calculated their average to obtain <η_s_> here. Mutational signatures of cultured human cells in which individual MMR genes, *msh2* or *msh6*, are knocked out (KO) using CRISPR ^22^ showed similar relations to <η_s_> (Fig. 5b) and so did the mutational signatures of *C. elegans* with individual KO of MMR genes, *pms2* or *mlh1* ^23^ (Fig. 5c). Limiting the comparison to T>A mutations that can arise through a TT or AA mismatch also showed a positive correlation between relative mutation frequency and <η_s_> (Fig. 5d for MMR deficient tumors and Fig. 5e for MMR KO human cells). Likewise, we observed a similar positive correlation for C>A & T>G mutations, both of which can occur through a CT or AG mismatch (Fig. 5f for MMO KO *C. elegans*). Overall, these analyses support a proposal that sequence dependence of MMR efficiency *in vivo*, as well as the context dependence for hypervariable CT/AG/TT mismatches is a characteristic conserved across different domains of life.

**Figure 5.**
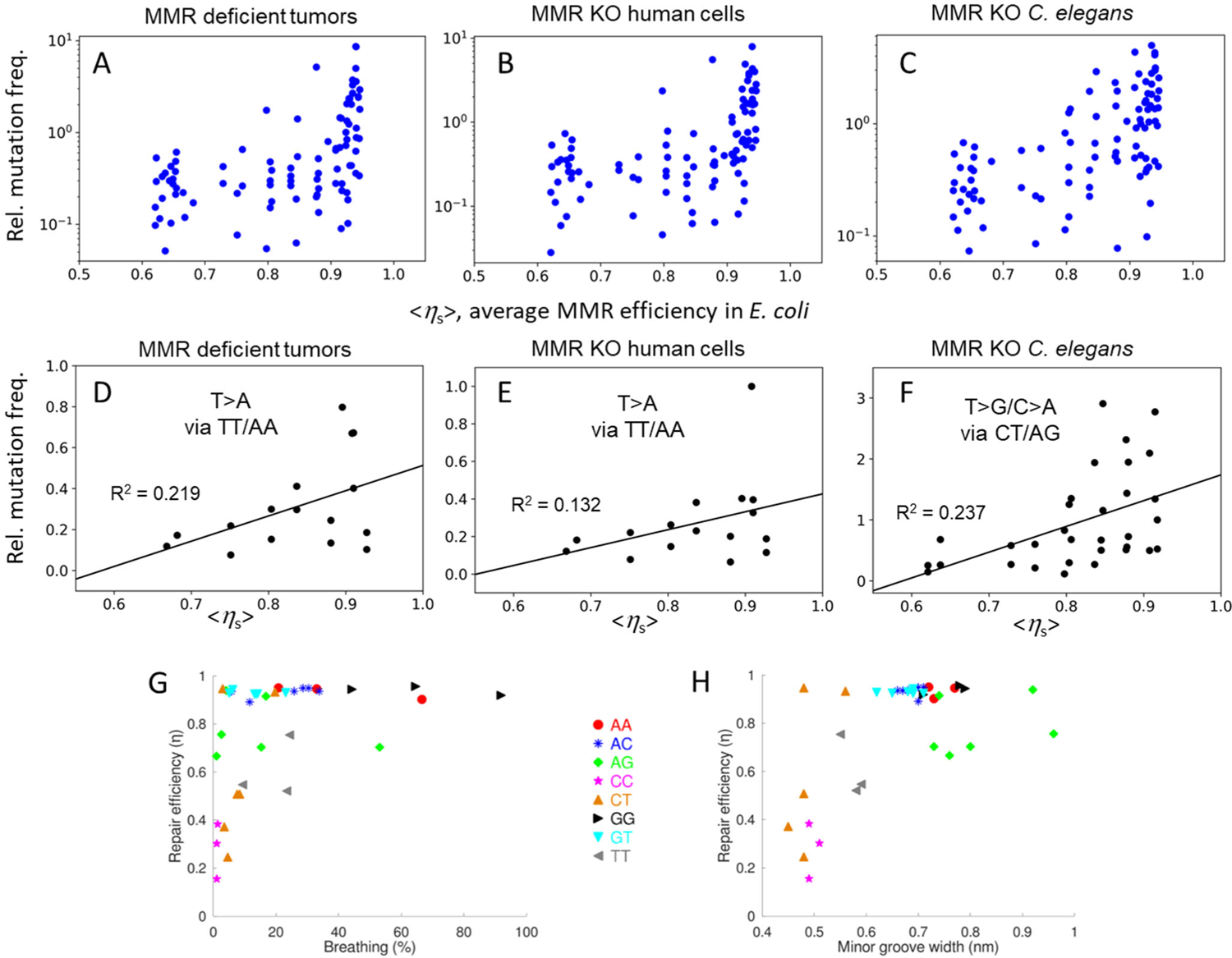
Sequence dependent mismatch repair conserved across species may arise from DNA physics. Relative mutation frequencies in trimer contexts vs MMR efficiency in *E. coli* averaged over mismatches that can give rise to the mutation, <*η*_s_>, for MMR deficient tumors (A), MMR KO human cells (B), and MMR KO *C. elegans* (C). The frequencies are normalized are against mutational signatures averaged over MMR proficient tumors. (D, E) Relative repair frequencies vs <*η*_s_> for T>A mutations that occur via TT/AA mismatches in MMR deficient tumors (D) and MMR KO human cells (E). (F) Relative repair frequencies vs <*η*_s_> for T>G and C>A mutations that occur via CT/AG mismatches. (G, H) Repair efficiency in *E. coli* vs base pair breathing percentage (G) and minor groove width (H) in MD simulations^24^.

### Poorly repaired mismatches share intrinsic physical properties

Rossetti *et al*. used all atom molecular dynamics (MD) simulations to analyze the physical properties of 13 bp dsDNA fragments containing a single mismatch, revealing how the mismatch types and different flanking sequences influence DNA structure dynamics^24^. We found that mismatches with low η_s_ values have narrower minor grooves according to MD simulations (Fig. 5g). Poorly repaired mismatches also have reduced local DNA dynamics such that the extent of base pair breathing is similar to that of fully base paired DNA (Fig. 5h). Overall, comparison to the existing data in the literature suggests that DNA physics is what distinguishes poorly repaired mismatched DNA from others. If so, the apparent conservation of sequence-dependence of *in vivo* MMR efficiencies, as revealed from comparison to mutational signatures in humans and *C. elegans*, can be potentially explained by the need to recognize diverse types and context of mismatches using the same arrangements of a set of amino acids contained in MutS and its homologues ^25^.

## Discussion

While the MMR efficiency was measured as early as 1980s, previous studies had limited throughput, sampling at most a dozen or so sequences^6,8-11,26^. We measured the *in vivo* repair efficiency of 5682 distinct mismatches in *sE. coli*, thereby sampling all possible mismatches within all pentameric sequence contexts at least once. The large dataset allowed us to discover selectively enhanced mutability through higher variable repair of TT mismatches and to account for context-dependent mutation frequencies in mismatch repair deficient tumors, cells, and animals.

High resolution structures have shown that a single base pair mismatch introduces only subtle structural changes in DNA only^27^, DNA bound to MutS_α_^25^, and DNA/RNA hybrid duplex bound to Cas9^28^. MutS_α_ recognizes diverse types of mismatches using the same set of amino acids and interactions^25^, and such a requirement may have resulted in a partial compromise where certain mismatches in some sequence contexts are poorly repaired. Interestingly, the context-dependent repair efficiency of TT mismatches appears to have been optimized to allow enhanced mutability when a mutation does not cause a large perturbation, for example only causing a synonymous codon change.

Because the mismatch-containing plasmid is not methylated at the GATC sites before its introduction into the cell, we did not expect to see a bias as to which of the two strands is used as a template for mismatch repair. However, we generally observed that the variable strand is over-represented in the repaired clans (Fig. 1c, Fig. S9). Unidirectional replication of the plasmid^29^ and asymmetric distribution of GATC sites in the plasmid (Fig. S10) are likely to be responsible for the observed asymmetry (Supplementary Discussion).

Relating substitution mutation frequencies observed upon MMR inactivation ^21-23^ to the MMR efficiencies we measured here is not always straightforward. Aside from species-specific effects, a substitution mutation can occur through two different types of mismatches and each mismatch type can occur through two polymerase error types. To be overrepresented in MMR-deficient samples, the replication polymerase should generate an error that leads to the substitution and the MMR system should normally repair that error efficiently. Conversely, for substitutions that are underrepresented in MMR-deficient samples, either the originating error is rarely made, or the error is poorly repaired. Therefore, we can expect perfect correlation between the relative mutation frequencies and repair efficiency reported in Fig. 5 only if there is no inter-species difference and if the DNA polymerase makes all errors with equal frequency, which is unlikely. Therefore, we find it remarkable that we can nonetheless see significant correlations. Comprehensive characterizations of replication polymerase error frequencies as a function of sequence context will be needed for examining this relation in more detail.

## Acknowledgments

This study was supported by an NIH grant R35 GM 122569 to T.H. and a National Science Foundation, Graduate Research Fellowship Program grant DGE-1746891 and NIH grant T32 GM007445 (Biochemistry, Cellular and Molecular Biology Graduate program) to J.S.Z. We thank Prof. Winston Timp for generously allowing the usage of his MiSeq instrument, and Keith Weninger, Hashim Al-Hashimi and Raluca Gordan for helpful discussion. T.H. is an investigator with the Howard Hughes Medical Institute.

## Data access

The full raw sequence data obtained as part of this study can be accessed via the NCBI Sequence Read Archive (SRA) in *.fastq format, under the accession number PRJNA917048. The source code of the analysis toolkit can be accessed through the T.K.’s Gitlab page: https://gitlab.com/tuncK/mmrseq-codes. Unpublished results and a detailed stepwise description of the analyzed data can be found in T.K.’s dissertation (http://jhir.library.jhu.edu/handle/1774.2/63982). Additional data will be made available by T.H. upon request.

## Author contributions

T.K. conceptualized the idea; T.K. designed the workflow on nucleic acids with help from T.H.; T.K. and C.L. designed the experiments on microbiologicals with help from T.H. and prepared the specimen; T.K. analyzed the data with help from T.H.. T.K., T.H. and K.D.H. interpreted the biological implications of the raw data. K.D.H. postulated the potential sequence dependence of the clan substitution histograms; S.M. conducted the follow-up experiments using biologically derived sequences with help from T.K.; J.A.L expressed, purified, and labeled the MutS protein; J.S.Z. performed single molecule microscopy experiments, which were designed and analyzed with help from T.H.; T.K. and T.H. drafted the manuscript; J.S.Z., T.K. and T.H. collaboratively finalized the manuscript.

## Materials and methods

### Preparation of traceable vector library (tUNC19)

We prepared chemically competent cells using NEBTurbo strain (NEB, C2984I) with *E. coli* Mix&Go Transformation kit with Zymobroth following manufacturer’s instructions (Zymo, T3001), hereafter referred to as Home-Made Competent Cells (HMCC). We transformed 50 pg pUC19 (NEB, N3041S; Genbank L09137) into a 100 μl HMCC aliquot by gentle agitation and extracted pUC19 plasmids by miniprep after overnight incubation in LB medium (Omega, E.Z.N.A. Plasmid Mini Kit I, D6942) inoculated by a single colony of this transformation. We used this product as template in a PCR reaction where one of the primers has 25 random bases as tracking barcode (primer P2, denoted by Ns) in blocks of five separated by T’s. An 800 μl batch of the reaction contains 50 ng pUC19 as template, 240 pmol of each primer (P1 and P2) and Phusion 2X mastermix with HF buffer (NEB, M0531S). The PCR mix was split into 100 μl aliquots and subjected to 35 cycles of 10 s 98 °C denaturation, 30 s 71 °C primer annealing and 60 s 72 °C elongation phases preceded by additional initial denaturation at 98 °C for 30 s and followed by 72 °C final extension for 2 min. The product was purified via QIAquick PCR purification kit (Qiagen, 28104) with 4 ml buffer PB supplemented with 20 μl 3 M NaAc, otherwise according to manufacturer’s instructions. This procedure has a typical yield above 10 μg, on whose product we generate sticky ends and remove remaining pUC19 templates by a triple digest in 1X Cutsmart buffer with 8 μl SacI-HF (NEB, R3156S), 8 μl XhoI (NEB, R0146S) and 8 μl DpnI (NEB, R0176S) in 400 μl final volume. The reaction mixture was incubated for 1 hour at 37 °C followed by a 30 min heat inactivation at 65 °C. The barcoded vector to be ligated was purified with QIAquick PCR purification kit out of this digestion mix.

### Design of DNA library sequences

To sample all trimer and pentamer sequence contexts and labor- and cost-effectively, we chose the consensus sequences of our libraries as the shortest sequence containing all sequence *k*-mers where *k* is 3 or 5. Commonly attributed to De Bruijn, the most compressed sequence on an alphabet Σ of size ||Σ||=n containing all *k*-mers, B(*n, k*) is a cyclic sequence of length *n*^*k*^ characters or *n*^*k*^ + *k* − 1 characters-long, if the former circular sequence is linearized. For our specific case of sampling DNA base motifs, Σ = {A, C, G, T}, *n*=4. We chose to sample in two different constructions all sequence trimers (3ML) and all sequence pentamers (5MLx, x=1,2,3…13) for which *k* = 3 and *k* = 5, respectively.

We generated such De Bruijn sequences by concatenating Lyndon words in lexicographic order^30^. Briefly, we generated a list of all necklaces of length *n* that are non-periodic and concatenated the individual words into a super-string. Here the Cyclic DEB Sequence is a cyclic sequence containing all sequence *k*-mers, Word denotes the constituting Lyndon words and incrementation is defined assuming base order A, C, G, T for DNA bases. To obtain a linear sequence representing all *k*-mers (Linear DEB Sequence), we suffixed this circular sequence with the initial *k*-1 bases of the sequence.

Unlike the trimer or pentamer cases, the length of the De Bruijn sequence containing all heptamers is impractically long (4^7^+7-1=16 390 bp). For the partial heptamer libraries (SSLx, x=1,2,3,4), we sought a sequence that represents all sequence heptamers that contain in their middle only a subset of the shorter sequence motifs that showed a notable variability in repair efficiency, namely CAAAA, AAAAT, GAAAT pentamers. Each library element is redundantly represented by a 7-base long mapping barcode that is separated by at least 2 substitution mutations from each other. As there will be 16 heptamers per pentamer, the length of the sequence obtained by naive concatenation is 16·7·3=336 bases long. While algorithmically finding the theoretically shortest DNA sequence is possible by brute force search or dynamic programming, we instead opted to find an approximate solution instead due to resource demanding nature of the exact solution. We resorted to a greedy algorithm to reduce the length of this sequence down to 244 bases, which we then split into 4 sub-libraries, each covering 80 bases. We defined the distance between DNA motifs d(x,y) as the minimum number of bases that the length of *k*-mer x increases by, if sequence y is to immediately follow x, taking into account the most extensive compression possible given the overlap between the two sequences. As an example, if x=ATATAA and y=TAATTA, d(x,y) = 3 since x and y can be combined to z=ATATAATTA, z contains both x and y, and |z|−|x|=9-6=3. We then loop till all nodes are processed, joining the two nodes separated by the shortest distance and marking the nodes as processed at each iteration. The routine quits by reporting a compressed concatenated consequence by traveling through the path formed by the joint edges, which is a shortened sequence but is not necessarily the theoretically shortest sequence. Due to the limit on solid-phase synthesized oligos, we split this sequence into 4 sub-libraries (SSL1, SSL2, SSL3, SSL4).

### Library preparation using oligoarrays

For the trimer library, we purchased a 4,007-member custom designed oligo library consisting of 132 bases long oligos (Twist Biosciences, CA). Each oligo contains a primer binding site (Z6 and Z13) for amplifications, a XbaI and XhoI cut site followed by a 7-nucleotides long mapping barcode. We computationally generated a 66 base long consensus sequence representing all sequence triplets in the form of a third order De Bruijn sequence on a 4-letter alphabet. This property ensures that all nearest neighbor sequence contexts are represented once and only once, hence achieving the theoretically most efficient way of sampling. Each individual member of the library differs from this consensus sequence by one base, and the mapping between the 7 bp barcode (separated by at least 2 mutations from each other) and the expected mismatch position is known a priori as a lookup table. To ensure that the observed repair efficiency is not an artifact of mapping barcode difference between different library members, each individual mismatch is represented by multiple tracking barcodes (∼3 distinct mapping barcodes per each mismatch for 5MLx, ∼20 per mismatch for 3ML). Barcodes generating undesirable extra restriction sites were computationally discarded during the design stage to avoid truncated inserts.

For the pentamer library (5MLx), we applied the same protocol except that we purchased a DNA library comprising a total of 9,828 different oligos, each 142 bases long (Genscript, NJ). The De Bruijn sequence sampling all pentamers is 1,028 bases long, and it not possible to procure a diverse DNA library consisting of oligos of this length with the current technology. We split the sequence down to 13 sub-sequences, sampling 84 out of 1028 bases with 4 base overlap at the termini. To encode the position and the type of the mismatch, we included 6-base long mapping barcodes, each separated by at least 2 mutations in the Hamming space. By design, the library consists of 13 sub-libraries and each sub-library is flanked by a different adapter pair combination for selective amplification of the chosen sub-library. By choosing the proper primer pair during the PCR (Z13/Z14 and Z1/Z2/Z3/Z4/Z5/Z6/Z7), we can obtain the sub-library of choice and hence each 5MLx sub-library was prepared, transformed and analyzed separately in the workflow. We purchased the sub-sampling heptamer libraries (SSLx) from Agilent (Wilmington, DE, G7220A) and applied the same protocol, except that we could obtain a clean product without making use of emulsion PCR (see **Mismatch library generation**), we hence omitted this step. We included all four sub-libraries in the same oligo pool and amplified by choosing the proper primer pair during the PCR (Z13 and Z1/Z2/Z3/Z4). The lists of all three oligo pools can be accessed among the Supplementary Materials.

For the measurements across a biologically relevant DNA sequence, we focused on *E. coli* RpoH encoding RNA polymerase sigma H (or sigma 32), as this sequence permutation is natively known to the cells used in our study as opposed to algorithmically generated sequences described above. To be able to assess whether competitive binding by other cytoplasmic proteins acts as a significant inhibitor of the repair response, we investigated across the promoter, 5’ UTR and N-terminal portion of the gene’s coding sequence (CDS). Due to the limitations on the maximum oligo length achievable in practice, we split the region of interest into 3 overlapping chunks of 170 nt. In this arrangement, RpoH1 covers the upstream regulatory region ([3601030, 3600861] of *E. coli* K12 genome U00096.3, minus strand), RpoH3 contains only CDS ([3600870,3600541]) whereas RpoH2 is the transition between these two and includes the start codon ([3600870,3600701]). The variable sequences were purchased from Agilent, each 170 nt long library member carrying a 7 nt long mapping barcode. The 186 nt long constant strands of the library were purchased as ultramers (IDT).

### Mismatch library generation

We amplified the oligo pool we received through two consecutive rounds of emulsion PCR (emPCR) following manufacturer’s instructions (ChimerX, catalog# 3600). Briefly, per each 50 μl aqueous reaction volume and 300 μl oil phase, we amplified 10 fmol of the original oligo library with unmodified 500 nM primer Z6 and 500 nM primer Z13 with Phusion polymerase (NEB, M0530L). The product was purified following manufacturer’s specifications, which comprise breaking up the emulsion in n-butanol, centrifugation for phase separation and DNA purification using a silicate column from the non-organic phase, and recovery of the amplified DNA with 50 μl aqueous elution buffer. We re-amplified this purified product with 500 nM Thio-Z13 and 500 nM Phospho-Z6 primers with Phusion polymerase. While we again employed emulsion conditions for 3ML for this second step, we could re-amplify 5MLx or SSLx without emulsion conditions. While highly dependent on the sequence template, a typical PCR reaction provided about 10 μg of product per ml of reaction volume after this second step. The thermal cycler protocol for both steps is 98 °C 30 s; 25x (98 °C 10s; 63 °C 20s; 72 °C 10s); 72 °C 2min; and the product was stored at 4 °C until further use.

We digested the phosphorylated strand by incubating each μg of the above obtained PCR product with 2.5U λ-exonuclease (NEB, M0262L) in 100 μl 1X reaction buffer at 37 °C for 1 hour followed by a 30 min heat-inactivation step at 75 °C. The ssDNA product was purified with a PCR purification column (Qiagen) and was thermally annealed to the constant strand (IDT) in T50 by slow cooling on a thermal cycler from 98 °C to 37 °C over 1 h, typically around 2 μM concentration. About 15 pmol of this partial duplex was extended with 1.25 U Taq polymerase (NEB, M0273L) in standard Taq buffer supplemented with 200 μM dNTP by incubation at 65 °C for 20 min. The product was purified using a PCR clean-up column (QIAGEN), and the elute was subjected to double digestion by 40 U SacI-HF (NEB, R3156L) and 40 U XhoI (NEB, R0146L) in 1X Cutsmart buffer for 2 h at 37 °C. The product was purified with a PCR clean-up column, and this elute is hereafter referred to as insert.

### Electroporation

We ligated the above insert with a tracking-barcoded vector library (tUNC19). A typical ligation reaction included about 1 - 2 μg vector, 3:1 insert:vector ratio, 150 kU T7 ligase (M0318L) in 1 ml 1X T7 DNA ligase reaction buffer incubated for 30 min at room temperature. The product was cleaned with a PCR clean-up column and eluted with 250 μl water. We mixed 5 μl of this elute with 100 μl ice-cold electrocompetent cells in a 1.5 ml tube and immediately transferred the cell-DNA mixture into a pre-chilled electroporation cuvette with a 1 mm gap width (Sigma-Aldrich, Z706078) and applied 1,700 V for about 5 ms (Eppendorf Eporator, 4309000027). We quickly washed the cuvette with 500 μl SOC growth medium twice and cells were allowed to recover at 37 °C for 1 hour in a 50 ml conical tube (Corning, CLS430829). We then added 9 ml room-temperature LB, with a final concentration of 100 μg/ml ampicillin and incubated overnight with constant 250 rpm shaking at 37 °C. We spread 100 μl of this final culture on an LB-agar plate with 100 μg/ml ampicillin to confirm efficient ligation and transformation. A typical experiment yields around 50-100 colonies after overnight incubation at 37°C, each colony corresponding to 100 expected clans due to the dilution factor.

### Next-generation sequencing

We extracted the plasmid library from a 10 ml overnight LB culture using a standard Miniprep kit following the prescribed protocol (Omega E.Z.N.A. Plasmid Mini Ki I, D6942). We used 5 ng of this elute as PCR template to amplify out the portion of interest using 0.5 μM of S1 and S2 primers with Phusion 2X mastermix. We employed 20 cycles of 10 s 98 °C denaturation, 20 s 63°C primer annealing 10s at 72 °C elongation phases preceded by additional initial denaturation at 98 °C for 30 s and followed by 72 °C final extension for 2 min. To clean up the product, we incubated the product mixture with 20 μl Ampure XP beads (Beckman Coulter, A63880) for 5 min. We retained the bead-bound material after incubating for 2 minutes on a magnetic rack (GE, 1201Q46). We washed the beads twice with 200 μl 80 % ethanol and eluted the material in 53 μl 10mM Tris pH 8.5 by incubation for 2 min. We collected about 45-50 μl bead-free liquid 2 min after placing the material on magnetic rack. We performed 8 additional cycles of PCR with Nextera 24-Index kit for indexing before sample pooling (Illumina, FC-121-1011), for which we used 7.5 μl of the elute as template, 7.5 μl of suitable i5 and i7 primers with 38 μl Phusion 2X. We followed the manufacturer’s recommended thermal cycling protocol (95 °C 3 min, 98 °C 30 s, 55 °C 30 s, 72 °C 30 s, 72 °C 5 min). We also bead-purified 56 μl of this final product with 56 μl Ampure-XP and eluted with 28 μl 10 mM Tris, pH 8.5 buffer, following the same procedure described above otherwise. We pooled the final products based on their Nanodrop and/or Qubit reading to the desired relative number density. While varying from sample to sample, we found a mixture of 480 μl Hbf buffer, 100-120 μl pooled-denatured 20 pM library and 10-20 μl 20 pM PhiX control library (Illumina, FC-110-3001) to provide a reasonable spot density in general. We typically used 300 cycles MiSeq v2 micro reagent kit (Illumina, MS-102-1002) to perform a paired-end sequencing for about 135-140 cycles each. For deeper sequencing, we used paired end 2×150 cycles sequencing on a Illumina HiSeq platform with 15 % PhiX spiked in (Genewiz, NJ).

### Calculation of repair efficiency

We retrieved raw *.fastq output from the MiSeq/HiSeq system and parsed with a custom C++ program and subsequently in GNU Octave v5.2. Processing a typical MiSeq dataset containing a few hundred thousand reads with this procedure takes around 5 minutes on a standard quadcore desktop computer. For samples obtained by HiSeq, we typically obtained around 50 million paired-end reads per sample, processing of which requires up to 25 GB RAM and 10-20 hours of CPU wall time on a 24-core node at Maryland Advanced Research Computing Center (MARCC), parallelized via OpenMP.

The major steps during analysis workflow are as follows: First, all the data is imported DNA by DNA, while reads are parsed to locate the constant adapter segments using a Needleman-Wunsch algorithm with gap and mismatch penalties of −1 and match gain of +1. We take the reverse complement of the paired end reads and add to the dataset to enhance the SNR by reducing the effect of sequencing errors. Using the *a priori* known library prototype, we extract the segment corresponding to the tracking barcode that immediately follows the end position of the adapter and the mismatch carrying library segment which follows the barcode after a 6 bp gap due to the SacI restriction site (GAGCTC). If this extracted library deviates from the library prototype by more than 5 substitutions, we attempt to re-align the sequence by Needleman-Wunsch algorithm. Reads that still deviate by more than 5 bases or do not carry a clearly identifiable adapter sequence are omitted. We export the barcode-read pairs as a text file (*.csf).

Second, we generate a unique set of all detected barcodes on a red-black tree, where exact duplicates are detected and recorded. To account for sequencing errors that could artificially diversify barcodes from the same clan, we introduced error tolerance by implementing a density-based scanning on the set of all detected barcodes, where the minimum density threshold of N=10 different reads should be reachable within ϵ=3 Hamming distance. To reduce memory usage, only a list of neighbors is stored rather than an explicit matrix listing pairwise distances. After all barcodes are processed, barcodes failing the density criterion (noise) are discarded, and the core-points together with all neighbors are reported as clans. Interested readers are referred to Ester et al^16^ for the description of this DBSCAN algorithm.

Third, the base distribution frequency in each clan is evaluated individually, as a 4x library length matrix per clan. For the purposes of the further analyses, each such matrix is considered as one data point. The output for each sample is hence a 3D tensor, which is output into a text file obeying the *.mat file format (*.hist).

Fourth, we import this *.hist file to GNU Octave v5.2 to interpret the results for each mismatch in the input library individually. To attribute a particular clan to the mismatch that its ancestor plasmid was carrying, we make use of the mapping barcodes. We use the maximum voted base combination in each clan to check a pre-determined lookup table. We omit clans whose mapping barcodes do not have an exact hit. Finally, we process the list of 4x length matrices clan by clan to count the number of unrepaired clans (U_ij_) or repaired clans using the constant (C_ij_) or the variable strand (V_ij_) as correct information source by the cell. Knowing these three variables for each mismatch position i and substituted base j, we can calculate both the repair efficiency (η_ij_) and strand choice bias (β_ij_) by:

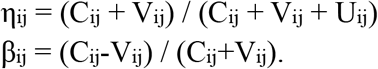

To distinguish among these three scenarios (U, C and V), we evaluate the proportion of reads constituting the clan that carry the substitution predicted by the barcode (s). Clans that were repaired using the constant strand (C) parallel the consensus sequence, and hence most reads will be unsubstituted (s≈0). But a repair keeping the variable strand (V) would be dominated by substituted reads (s≈1); unrepaired clans (U) are expected to follow a distribution around intermediate s’s. For the C vs. U decision problem, we implemented a system-wide ad hoc fixed threshold of c_low_(i) = 0.1.

To compensate for the shift in the peak position from s=1 towards s=0.8 along the mismatch library, we applied a moving threshold to decide between U and V cases. To find the high-cutoff *c*_high_(*i*) which depends on the mismatch position *i*, we used curve fitting on the substitution histogram of each individual mismatch using the function below:

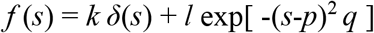

where *k, l, p* and *q* are the parameters to optimize the fit of the model function to the observed histogram. We performed an interpolation by fitting all p_ij_ with a 6^th^ degree polynomial to find an interpolation for the peak position along *s*, <*p*_i_>_j_ that depends on substitution position *i*, but not on the substituted base type *j*. Ad hoc, we considered all clans with a substitution frequency above *c*_high_ = <*p*_i_>_j_ − 0.15 as repaired by keeping the variable strand of the library.

### Preparation of fluorescently labeled MutS

*E. coli* MutS was cloned with a N-terminal hexa-histidine and sortase (LPETG) tag into a pET expression plasmid. MutS was expressed in BL21-AI cells with 1 L of culture. The culture was grown at 37 °C until it reached an OD_600_ of 0.3. The temperature was lowered to 16 °C and 0.2 % L-(+)-Arabinose was added. After 30 minutes 0.05 mM IPTG was added, and the culture grew for 16 hrs. The cells were collected and resuspend in Buffer A (25 mM HEPES pH 7.8, 10 % glycerol, 0.8 μg/mL pepstatin, 1 μg/mL pepstatin and 87 μg/mL PMSF) containing 300 mM NaCl and 20 mM imidazole. The resuspended cell pellet was flash frozen and stored at –80 °C. For the purification of MutS, all steps were carried out at 4 °C. The cell pellet was thawed and lysed by sonication. The lysate was clarified by centrifuging at 120,000 x g for 1 h at 4 °C. The supernatant was then applied to a 3 mL Ni-NTA (Qiagen) column. The Ni-NTA column was washed with 10 column volumes of Buffer A containing 800 mM NaCl and 20 mM imidazole followed by 5 column volumes of Buffer A containing 90 mM NaCl and 20 mM imidazole. The Ni-NTA column was then eluted directly onto a tandem 3 mL Heparin-Sepharose (Cytiva) column with 5 column volumes of Buffer A containing 90 mM NaCl and 200 mM imidazole, washed with 10 column volumes of Buffer B (25 mM HEPES pH 7.8 and 10 % glycerol) containing 100 mM NaCl and eluted with a 20 mL linear gradient of Buffer B containing 100 mM to 1M NaCl. Peak fractions were analyzed by PAGE and combined. The combined MutS fractions were fluorescently labeled using a sortase transpeptidase. A total of 13 nmol of MutS was combined with 52 nmol of purified *S. aureus* sortase 5M, and 130 nmol of purified Cy3-CLPETGG labeled peptide. The labeling reaction was carried out at 4 °C for 1 hr in Buffer B containing 10 mM CaCl_2_ and 300 mM NaCl. The reaction was then stopped by the addition of 20 mM EGTA. The excess label was removed by passing the reaction mix through a 10K MWCO Zeba spin desalting column (Thermo Scientific) followed by application onto a 1 mL Heparin-Sepharose column. Bound material was step eluted with Buffer C (25 mM HEPES pH 7.8, 600 mM NaCl, 10 % glycerol, 0.1 mM EDTA, 1 mM DTT, 0.8 μg/mL pepstatin, 1 μg/mL pepstatin and 87 μg/mL PMSF), which removed the Sortase protein and concentrated the MutS. Peak fractions were dialyzed into storage buffer (25 mM HEPES pH 7.8, 150 mM NaCl, 20 % glycerol, 0.1 mM EDTA and 1 mM DTT), flash frozen and stored at −80 °C. We determined a 48 % MutS monomer labeling efficiency.

### DNA substrates for *in vitro* FRET experiments

PAGE or HPLC purified oligodeoxynucleotides, modified with either a 5’ biotin or amino-C6 dT, were purchased from Integrated DNA Technologies Inc (Supplemental Table 1). The amine modified dT nucleotides were labeled with Cy5 in a 20 µL reaction containing 10 mM (200 nmol) NHS-Cy5 (GE Healthcare, PA15100), 0.1 mM (2 nmol) amine modified oligo, 0.1 M freshly prepared sodium bicarbonate, and 0.2 M DMSO. The reaction was incubated at 40 °C overnight. Free dye was removed from the sample using ethanol precipitation. Briefly, threefold volume of cold 100 % ethanol was added to the sample and placed at –80 °C for two hours. The sample was centrifuged at 15,000 rpm, at 4 °C for 40 min. The resulting pellet was washed several times with cold 70 % ethanol and then placed in a dark drawer and allowed to air dry. The labeled DNA oligo was then resuspended in 10 µl TE buffer (10 mM Tris and 1 mM EDTA, pH 8.0).) DNA substrates were prepared by annealing the paired top and bottom oligos in an equal molar ratio, 1 µM each. The solution was incubated at 98 °C for 2 min before slowly cooling down until 4 °C over one hour. The annealed DNA substrates were stored at –20 °C.

### Quartz slide passivation

Quartz slides and glass coverslips were passivated with polyethylene glycol [98 % polyethylene glycol (PEG), 2 % biotin-PEG, JHU Slide Production Core] using previously established methods^31^. Once passivated, quartz slides and coverslips were assembled into flow chambers with double-sided tape and epoxy.

### In vitro smFRET MutS binding reaction conditions

Fluorescently labeled DNA were immobilized in imaging chambers assembled from PEG-passivated quartz slides. After assembly, the chamber was washed with 10 mM Tris-HCl (pH 8.0), 50 mM NaCl and 0.1 mg/mL BSA. 40 pM biotinylated and Cy5 labeled double strand DNA (in 10 mM Tris-HCl (pH 8.0), 50 mM NaCl and 0.1 mg/mL BSA (NEB, B9000S)) was incubated in the chamber, in which a quartz slide functionalized with PEG and PEG-biotin was coated with neutravidin (Thermo scientific, 31000), for approximately five minutes to allow the biotinylated DNA to bind a neutravidin immobilized on the slide. The chamber was washed with 10 mM Tris-HCl (pH 8.0), 50 mM NaCl and 0.1 mg/mL BSA and then subsequently washed out with 40-fold excess volume blocking buffer (20 mM Tris-HCl, pH 7.5, 2 mM EDTA, 50 mM NaCl, 0.0025% (v/v) Tween 20, 0.1 mg/ml BSA) to remove free DNA molecules from the chamber. The tethered DNA was always imaged before an experiment to validate the DNA density on the slide was appropriate. To initiate the MutS binding studies, 5 nM Cy3 labeled *E. coli* MutS with 1 mM ATP in binding buffer (30 mM Tris-HCl, pH 7.5, 5 mM MgCl_2_, 100 mM potassium glutamate, 0.1 mM DTT, 20 mg/mL Acetylated BSA (Invitrogen, AM2614) and 0.0025 % Tween 20 (BioRad, 170-6531)) supplemented with oxygen-scavenging system (∼3 mM Trolox (Arcos organics, 53188-071), 0.8 % (w/v) dextrose monohydrate (Sigma Aldrich, D9559-1KG), 2.5 mM PCA (3,4-Dihydroxybenzoic acid) (Sigma, 37580-25G-F), and 50 nM PCD (rPCO (recombinant protocatechuate 3,4-Dioxygenase from bacteria) (OYC, 46852004)), to minimize fluorophore photobleaching, was added to the chamber and imaged. All single molecule experiments were performed at room temperature.

The fluorescent microscopy signals were collected with a time resolution and exposure t of 50 ms, and the FRET traces shown were obtained using 10 frames of Cy5 excitation, followed by 1480 frames of Cy3 excitation followed by 10 frames Cy5 excitation.

### smFRET microscope instrumentation

Total internal reflection fluorescence (TIRF) microscopy was performed on a Nikon Eclipse Ti microscope equipped with a Nikon perfect focus system and a homebuilt prism-TIRF module^31^. The system was driven by homebuilt software. A Nikon 60×/1.27 NA (numerical aperture) objective (CFI Plan Apo IR 60XC WI) was used. Illumination was provided by solid-state lasers (Coherent, 641 nm; and Shanghai Dream Lasers Technology, 543). Emission was collected using long-pass filters (T540LPXR UF3, T635LPXR UF3, and T760LPXR UF3) and a custom laser-blocking notch filter (ZET488/543/638/750M) from Chroma. Images were recorded using an electron-multiplying charge-coupled device (Andor iXon 897).

### FRET data analysis

Raw movies were processed into raw fluorescence intensity time traces using custom written IDT scripts and analyzed using a custom written MATLAB script. Dwell times were collected by manual inspection using custom MATLAB scripts. *τ*_-1_ are the dwell times of the high FRET state when Cy3 labeled MutS binds on the Cy5 labeled mismatched DNA, but there is no low FRET state after the initial high FRET state. *τ*_2_ are the dwell times of the high FRET state when Cy3 labeled MutS binds on the Cy5 labeled mismatched DNA and is followed by a low FRET state.

To determine *k*_-1_ and *k*_2_, the *τ*_-1_ and *τ*_2_ dwell times values from 54 traces per film for three films (where each film was imaged in a separate area/region/location of the slide) were plotted in Origin (OriginLab) to generate a histogram of the dwell times. The bin sizes (0.05 or 0.1 seconds) were manually chosen to best fit the data. The histograms were then fitted to a single exponential decay curve to obtain the rate constant. The single exponential decay function used:

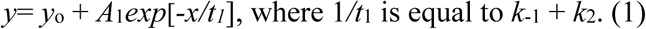

From these selected traces, the number of binding events where MutS initially bound to the mismatch from a high FRET state followed by a low FRET state were counted and divided by the number of binding events where MutS initially bound to the mismatch from a high FRET state, but the event was not followed by a low FRET state. This branching ratio (*N*_2_/*N*_-1_) was set equal to the ratio of the rate constants *k*_-1_/*k*_2_:

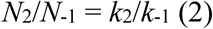

Equations (1) and (2) were solved in order to obtain *k*_-1_ and *k*_2_, resulting in *k*_-1_=(1/*t*_1_)/(1+ (*N*_2_/*N*_1_) and *k*_2_= (*N*_2_/*N*_-1_)\**k*_-1_.

To determine *k*_1_ (the rate at which MutS binds specifically to a mismatch), a subset of traces was analyzed where the Cy5 labeled DNA was still fluorescent at the end of the long films/traces. Any traces where Cy5 had photobleached before the end of the film were not included in the data set used to calculate *k*_1_. For each trace, the total number of binding events, where the Cy3 labeled MutS initially binds to the Cy5 labeled mismatched DNA resulting in a high FRET state (regardless of whether a low FRET state followed the high FRET state or not) were calculated and averaged over the total number of traces used in the data set. This average number of high FRET binding events was then divided by the total time of green excitation during the long trace (74 seconds), resulting in the *k*_1_ value. For a given experiment this was done for 3 films taken at different regions of the slide. All experiments were performed in triplicates and the resulting error bars are the standard errors from the triplicate experiments.

