## Supplementary Materials for "Massively parallel single molecule tracking of sequence-dependent DNA mismatch repair *in vivo*"

Supplementary Table 1

| DNA Construct | Annealed DNA Strand sequences (5' to 3') |
|---|---|
| No Mismatch | 5'-/5Biosg/TTGGGCGCCCGGACGAGCACCAGAACAAAGGTCGTAGTGCTCCTACTGATCATAATGTTCTTATTT-3' |
| | 5'- AAATAAGAACATTATGATCAGTAGGAGCAC /iAmMC6T/ ACGACCTTTGTTCTGGTGCTCGTCCGGGCGCCCAA -3' |
| GT Mismatch | 5'-/5Biosg/TTGGGCGCCCGGACGAGCACCAGAACAAGGGTCGTAGTGCTCCTACTGATCATAATGTTCTTATTT-'3 |
| | 5'- AAATAAGAACATTATGATCAGTAGGAGCAC /iAmMC6T/ ACGACCTTTGTTCTGGTGCTCGTCCGGGCGCCCAA -3' |
| CC Mismatch | 5'-/5Biosg/TTGGGCGCCCGGACGAGCACCAGAACAAACGTCGTAGTGCTCCTACTGATCATAATGTTCTTATTT-'3 |
| | 5'- AAATAAGAACATTATGATCAGTAGGAGCAC /iAmMC6T/ ACGACCTTTGTTCTGGTGCTCGTCCGGGCGCCCAA -3' |
| TT Mismatch | 5'-/5Biosg/TTGGGCGCCCGGACGAGCACCAGAACAATGGTCGTAGTGCTCCTACTGATCATAATGTTCTTATTT-'3 |
| | 5'- AAATAAGAACATTATGATCAGTAGGAGCAC /iAmMC6T/ ACGACCTTTGTTCTGGTGCTCGTCCGGGCGCCCAA -3' |
| AG Mismatch | 5'-/5Biosg/TTGGGCGCCCGGACGAGCACCAGAACAAAGGgCGTAGTGCTCCTACTGATCATAATGTTCTTATTT-3' |
| | 5'- AAATAAGAACATTATGATCAGTAGGAGCAC /iAmMC6T/ ACGACCTTTGTTCTGGTGCTCGTCCGGGCGCCCAA -3' |
| AC Mismatch | 5'-/5Biosg/TTGGGCGCCCGGACGAGCACCAGAACAAAGGcCGTAGTGCTCCTACTGATCATAATGTTCTTATTT-3' |
| | 5'- AAATAAGAACATTATGATCAGTAGGAGCAC /iAmMC6T/ ACGACCTTTGTTCTGGTGCTCGTCCGGGCGCCCAA -3' |

| | |
|---|---|
| AA Mismatch | 5'-/5Biosg/TTGGGCGCCCGGACGAGCACCAGAACAAAGGaCGTAGTGCTCCTACTGATCATAATGTTCTTATTT-3' |
| | 5'- AAATAAGAACATTATGATCAGTAGGAGCAC /iAmMC6T/ ACGACCTTTGTTCTGGTGCTCGTCCGGGCGCCCAA -3' |

**List of DNA oligos used for MMR-seq**

All oligos were purchased from IDT. Primers were standard desalted, whereas oligos used for the construction of the mismatch libraries were bought with HPLC purification (C-strands).

Primer P1
/5/ TTTTTTT CTCGAG GCAAGCTTGGCGTAATCATGGTCAT /3/

Primer P2 - 25bp barcode
/5/ TTTTTTT GAGCTC NNNNNTNNNNNTNNNNNTNNNNNTNNNNN TGCGGTATTTCA-CACCGCATATGGT /3/

Primer P2 - 20bp barcode
/5/ TTTTTTT GAGCTC NNNNNTNNNNNTNNNNNTNNNNN TGCGGTATTTCACACCG-CATATGGT /3/

Primer P4
/5/ TTTTTTT CTCGAG GGTGCCTAATGAGTGAGCTAACTCAC /3/

Primer inv P1
/5/ TTTTTTT GAGCTC GCAAGCTTGGCGTAATCATGGTCAT /3/

Primer inv P2
/5/ TTTTTTT CTCGAG NNNNNTNNNNNTNNNNNTNNNNNTNNNNN TGCGGTATTTCA-CACCGCATATGGT /3/

pUC19 Sanger sequencing primer
/5/ CACAGCTTGTCTGTAAGCGG /3/

Primer S1
/5/ TCGTCGGCAGCGTCAGATGTGTATAAGAGACAG CATATGCGGTGTGAAATACCGCA /3/

Primer S2
/5/ GTCTCGTGGGCTCGGAGATGTGTATAAGAGACAG GCTAGTACCTCAATATAGACTC-CCT /3/

Primer S2_2
/5/ GTCTCGTGGGCTCGGAGATGTGTATAAGAGACAG AAGCTTGCCTCGACAGAATAG-GAAC /3/

Primer S2_3
/5/ GTCTCGTGGGCTCGGAGATGTGTATAAGAGACAG GCCAAGCTTGCCTCGAGCT /3/

Primer S2_4
/5/ GTCTCGTGGGCTCGGAGATGTGTATAAGAGACAG ATTACGCCAAGCTTGCCTCGAG /3/

Primer S2_5
/5/ GTCTCGTGGGCTCGGAGATGTGTATAAGAGACAG ATTACGCCAAGCTTGCCTCGA /3/

Primer S2_6
/5/ GTCTCGTGGGCTCGGAGATGTGTATAAGAGACAG ATTACGCCAAGCTTGCGAGCT /3/

Primer Phospho-Z1
/5/5Phos/ ATGGGACCGCATCGTAGCTT /3/

Primer Phospho-Z2
/5/5Phos/ CGGCTGAATGGTACCCGATA /3/

Primer Phospho-Z3
/5/5Phos/ GAGCGCAGCTGGTGTAGACA /3/

Primer Phospho-Z4
/5/5Phos/ CGACTTCGAATTCAGCACGT /3/

Primer Phospho-Z5
/5/5Phos/ ACACCCGCTCGATCCCTTAT /3/

Primer Phospho-Z6
/5/5Phos/ GGTTGAACAACCGCCGGTAT /3/

Primer Phospho-Z7
/5/5Phos/ AAATTGCCGTTGCGGATTTC /3/

Primer Thio-Z13, (*) indicates position of a phosphothioate bond
/5/biotin/ C*G*C*T*C*CTTGTTGTACTCGCA /3/

Primer Thio-Z14
/5/biotin/ T*G*A*C*G*GAGGATAGAAGGCCA /3/

Library sequence prototypes. Adaptor sites are indicated in blue and red, while the restriction sites are shown in bold font. N's indicate the random bases that constitute the mapping barcodes, ideally containing ACGT bases with 25% probability each. Required primer pair for amplification are indicated in parentheses following the library name.

3ML1 prototype (Z13, Z6)
/5/ TTCTAGA CGCTCCTTGTTGTACTCGCA GAGCTC NNNNNNN
TTTATTCTTGTAATACTAGTCATCCTCGTGATGCTGGAAACAAGACCACGAGCAGGCCCGCG
GGTT CTCGAG ATACCGGCGGTTGTTCAACC /3/

5ML1 prototype (Z13, Z1)
/5/ CGCTCCTTGTTGTACTCGCA GAGCTC NNNNNN
AAAAACAAAAGAAAATAAACCAAACGAAACTAAAGCAAAGGAAAGTAAATCAAATGAAA
TTAACACAACAGAACATAACCCAAC CTCGAG AAGCTACGATGCGGTCCCAT /3/

5ML2 prototype (Z13, Z2)
/5/ CGCTCCTTGTTGTACTCGCA GAGCTC NNNNNN
CAACCGAACCTAACGCAACGGAACGTAACTCAACTGAACTTAAGACAAGAGAAGATAAGC
CAAGCGAAGCTAAGGCAAGGGAAG CTCGAG TATCGGGTACCATTCAGCCG /3/

5ML3 prototype (Z13, Z3)
/5/ CGCTCCTTGTTGTACTCGCA GAGCTC NNNNNN
GAAGGTAAGTCAAGTGAAGTTAATACAATAGAATATAATCCAATCGAATCTAATGCAATGGA
ATGTAATTCAATTGAATTTACA CTCGAG TGTCTACACCAGCTGCGCTC /3/

5ML4 prototype (Z13, Z4)
/5/ CGCTCCTTGTTGTACTCGCA GAGCTC NNNNNN
TACACCACACGACACTACAGCACAGGACA
CATCACATGACATTACCAGACCATACCCCACCCGACCCTACCGCACCGGACC CTCGAG
ACGTGCTGAATTCGAAGTCG /3/

5ML5 prototype (Z13, Z5)

/5/ CGCTCCTTGTTGTACTCGCA GAGCTC NNNNNN GACCGTACCTCACCTGACCTTACGAGACGATACGCCACGCGACGCTACGGCACGGGACGGT ACGTCACGTGACGTTACTAGACT CTCGAG ATAAGGGATCGAGCGGGTGT /3/

5ML6 prototype (Z13, Z6)
/5/ CGCTCCTTGTTGTACTCGCA GAGCTC NNNNNN GACTATACTCCACTCGACTCTACT-GCACTGGACTGTACTTCACTTGACTTTAGAGCAGAGGAGAGTAGATCAGATGAGATTAGC CTCGAG ATACCGGCGGTTGTTCAACC /3/
5ML7 prototype (Z13, Z7)
/5/ CGCTCCTTGTTGTACTCGCA GAGCTC NNNNNN TAGCATAGCCCAGCCGAGCCTAGCGCAGCGGAGCGTAGCTCAGCTGAGCTTAGGATAGGCC AGGCGAGGCTAGGGCAGGGGAGG CTCGAG GAAATCCGCAACGGCAATTT /3/

5ML8 prototype (Z14, Z1)
/5/ TGACGGAGGATAGAAGGCCA GAGCTC NNNNNN GAGGGTAGGTCAGGTGAGGT TAGTATAGTCCAGTCGAGTCTAGTGCAGTGGAGTGTAGTTCAGTTGAGTTTATATCATATGAT A CTCGAG AAGCTACGATGCGGTCCCAT /3/

5ML9 prototype (Z14, Z2)
/5/ TGACGGAGGATAGAAGGCCA GAGCTC NNNNNN GATATTATCCCATCCGATCCTATCGCATCGGATCGTATCTCATCTGATCTTATGCCATGCGATG CTATGGCATGGGATGGTATG CTCGAG TATCGGGTACCATTCAGCCG /3/

5ML10 prototype (Z14, Z3)
/5/ TGACGGAGGATAGAAGGCCA GAGCTC NNNNNN TATGTCATGTGATGTTATTC-CATTCGATTCTATTGCATTGGATTGTATTTCATTTGATTTCCCCCGCCCCTCCCGGCCCGTCC CTCGAG TGTCTACACCAGCTGCGCTC /3/

5ML11 prototype (Z14, Z4)
/5/ TGACGGAGGATAGAAGGCCA GAGCTC NNNNNN GTCCCTGCCCTTCCGCGCCGCTC CGGGCCGGTCCGTGCCGTTCCTCGCCTCTCCTGGCCTGTCCTTGCCTTTCGCGGCGCGTCG CTCGAG ACGTGCTGAATTCGAAGTCG /3/

5ML12 prototype (Z14, Z5)
/5/ TGACGGAGGATAGAAGGCCA GAGCTC NNNNNN GTCGCTGCGCTTCGGCTCGGGGCGGGTC GTGCGGTTCGTCTCGTGGCGTGTCGTTGCGTTTCTCTGCTCTTCTGGGCTGGTCT CTCGAG ATAAGGGATCGAGCGGGTGT /3/

5ML13 prototype (Z14, Z6)

/5/ TGACGGAGGATAGAAGGCCA GAGCTC NNNNNN
GTCTGTGCTGTTCTTGGCTTGTCTTTGC
GTGTGGTTTGTGTTGTTTTAAAAACAAAAGAAAATAAAC CTCGAG
ATACCGGCGGTTGTTCA /3/

SSL1 prototype (Z13, Z1)
/5/ CGCTCCTTGTTGTACTCGCA GAGCTC NNNNNNN
GGTTTTCTTTAGGTTTTTAGCTTTAGCTTTATTTTATTTTACCTTTAAGGTTTTGTTTTATCCTT
TATTTTAGTTTTAAG CTCGAG AAGCTACGATGCGGTCCCAT /3/

SSL2 prototype (Z13, Z2)
/5/ CGCTCCTTGTTGTACTCGCA GAGCTC NNNNNNN
TTATTTTAGTTTTAAGTTTTGTTTTCTTTATCCTTTACCTTTAGCTTTAACTTTAGTTTTCTTTA
ACTTTTAGCTTTACT CTCGAG TATCGGGTACCATTCAGCCG /3/

SSL3 prototype (Z13, Z3)
/5/ CGCTCCTTGTTGTACTCGCA GAGCTC NNNNNNN
AACTTTTAGCTTTACTTTTATCGTTTTGTATCGTTTTCTTTACGTTTTACGTTTTTACTTTACTT
TAACTT CTCGAG TGTCTACACCAGCTGCGCTC /3/

SSL4 prototype (Z13, Z4)
/5/ CGCTCCTTGTTGTACTCGCA GAGCTC NNNNNNN
TACTTTACTTTAACTTTATTTTAACTTTTACTTTTAAGGTTTTAGTTTTTAAGGTTTTCTTTAGG
TTTTTAGCTTTAGCT CTCGAG ACGTGCTGAATTCGAAGTCG /3/

RpoH1 prototype
CGCTCCTTGTTGTACTCGCA GAGCTC NNNNNNN TAATAAAAGC GTGTTATACT
CTTTCCCTGC AATGGGTTCC GTAGCAGGGA AAGAGACCCC
GTTGTCTCTT CCCGGTATTT CATCTCTATG TCACATTTTG TGCGTAATTT ATTCACAAGC
TTGCATTGAA CTTGTGGATA AAATCACGGT CTGATAAAAC AGTGAATGAT
CTCGAG AAGCTACGATGCGGTCCCAT

RpoH2 prototype
CGCTCCTTGTTGTACTCGCA GAGCTC NNNNNNN AGTGAATGAT AACCTCGTTG
CTCTTAAGCT CTGGCACAGT TGTTGCTACC ACTGAAGCGC
CAGAAGATAT CGATTGAGAG GATTTGAATG ACTGACAAAA TGCAAAGTTT AGCTTTAGCC
CCAGTTGGCA ACCTGGATTC CTACATCCGG GCAGCTAACG CGTGGCCGAT CTCGAG
AAGCTACGATGCGGTCCCAT

RpoH3 prototype

CGCTCCTTGTTGTACTCGCA GAGCTC NNNNNNN CGTGGCCGAT GTTGTCGGCT
GACGAGGAGC GGGCGCTGGC TGAAAAGCTG CATTACCATG
GCGATCTGGA AGCAGCTAAA ACGCTGATCC TGTCTCACCT GCGGTTTGTT GTTCATATTG
CTCGTAATTA TGCGGGCTAT GGCCTGCCAC AGGCGGATTT GATTCAGGAA
CTCGAG AAGCTACGATGCGGTCCCAT

We annealed the ssDNA libraries we obtained to chemically synthesized oligos, 100 bases each, purchased with HPLC purification individually:

C-3ML1
/5/ CGGTAT CTCGAG
AACCCGCGGGCCTGCTCGTGGTCTTGTTTCCAGCATCACGAGGATGACTAGTATTACAAGA
ATAAA /3/

C-5ML1
/5/ ATCGTAGCTT CTCGAG GTTGGGTTATGTTCTGTTGTGTTAATTTCATTTGATT-
TACTTTCCTTTGCTTTAGTTTCGTTTGGTTTATTTTCTTTTGTTTTT /3/

C-5ML2
/5/ GTACCCGATA CTCGAG CTTCCCTTGCCTTAGCTTCGCTTGGCTTATCTTCTCTTGTCT-
TAAGTTCAGTTGAGTTACGTTCCGTTGCGTTAGGTTCGGTTG /3/

C-5ML3
/5/ GGTGTAGACA CTCGAG TGTAAATTCAATTGAATTACATTCCATTGCATTAGATTC-
GATTGGATTATATTCTATTGTATTAACTTCACTTGACTTACCTTC /3/

C-5ML4
/5/ TTCAGCACGT CTCGAG GGTCCGGTGCGGTAGGGTCGGGTGGGGTATGGTCTG-
GTAATGTCATGTGATGTACTGTCCTGTGCTGTAGTGTCGTGTGGTGTA /3/

C-5ML5
/5/ GATCCCTTAT CTCGAG AGTCTAGTAACGTCACGTGACGTACCGTCCCGTGCCG-
TAGCGTCGCGTGGCGTATCGTCTCGTAAGGTCAGGTGAGGTACGGTC /3/

C-5ML6
/5/ CCGCCGGTAT CTCGAG
GCTAATCTCATCTGATCTACTCTCCTCTGCTCTAAAGTCAAGTGAAGTACAGTCCAGTGCAG
TAGAGTCGAGTGGAGTATAGTC /3/

C-5ML7
/5/ TGCGGATTTC CTCGAG CCTCCCCTGCCCTAGCCTCGCCTGGCCTATCCTAAGCTCAGCT-

GAGCTACGCTCCGCTGCGCTAGGCTCGGCTGGGCTATGCTA /3/

C-5ML8
/5/ ATCGTAGCTT CTCGAG TATCATATGATATAAACTCAACTGAACTACACTCCACTG-CACTAGACTCGACTGGACTATACTAACCTCACCTGACCTACCCTC /3/

C-5ML9
/5/ GTACCCGATA CTCGAG CATACCATCCCATGCCATAGCATCGCATGGCATAAGATCA-GATGAGATACGATCCGATGCGATAGGATCGGATGGGATAATATC /3/

C-5ML10
/5/ GGTGTAGACA CTCGAG GGACGGGCCGGGAGGGGCGGGGGAAAATCAAATGAAAT-ACAATCCAATGCAATAGAATCGAATGGAATAACATCACATGACATA /3/

C-5ML11
/5/ TTCAGCACGT CTCGAG CGACGCGCCGCGAAAGGCAAGGACAGGCCAGGAGAG-GCGAGGAACGGCACGGACCGGCCCGGAGCGGCGCGGAAGGGCAGGGAC /3/

C-5ML12
/5/ GATCCCTTAT CTCGAG AGACCAGCCCAGAAGAGCAGAGAAACGCAACGACACGC-CACGAGACGAACCGCACCGACCCGCCCCGAGCCGAAGCGCAGCGAC /3/

C-5ML13
/5/ CCGCCGGTAT CTCGAG GTTTATTTTCTTTTGTTTTTAAAAACAACACAAACCA-CACCAACCCACCCCCAAAAGCAAAGACAAGCCAAGAACAGCACAGAC /3/

C-SSL1
/5/ CCGCATCGTAGCTT CTCGAG CTTAAAACTAAAATAAAGGATAAAACAAAACCT-TAAAGGTAAAATAAAATAAAGCTAAAGCTAAAAACCTAAAGAAAACC /3/

C-SSL2
/5/ AATGGTACCCGATA CTCGAG AGTAAAGCTAAAAGTTAAAGAAAACTAAAGTTAAAGC-TAAAGGTAAAGGATAAAGAAAACAAAACTTAAAACTAAAATAA /3/

C-SSL3
/5/ AGCTGGTGTAGACA CTCGAG AAGTTAAAGTAAAGTAAAAACGTAAAACGTAAA-GAAAACGATAAAAACAAAACAAAACGATAAAAGTAAAGCTAAAAGTT /3/

C-SSL4
/5/ CGAATTCAGCACGT CTCGAG AGCTAAAGCTAAAAACCTAAAGAAAACCTTAAAAAC-TAAAACCTTAAAAGTAAAAGTTAAAATAAAGTTAAAGTAAAGTA /3/

RpoH oligos:
C-H1
ATCGTAGCTT CTCGAG
ATCATTCACTGTTTTATCAGACCGTGATTTTATCCACAAGTTCAATGCAAGCTTGTGAATAAA
TTACGCACAAAATGTGACATAGAGATGAAATACCGGGAAGAGACAACGGGGTCTCTTTCCC
TGCTACGGAACCCATTGCAGGGAAAGAGTATAACACGCTTTTATTA

C-H2
GTACCCGATA CTCGAG
ATCGGCCACGCGTTAGCTGCCCGGATGTAGGAATCCAGGTTGCCAACTGGGGCTAAAGCTA
AACTTTGCATTTTGTCAGTCATTCAAATCCTCTCAATCGATATCTTCTGGCGCTTCAGTGGTA
GCAACAACTGTGCCAGAGCTTAAGAGCAACGAGGTTATCATTCACT

C-H3
GGTGTAGACA CTCGAG
TTCCTGAATCAAATCCGCCTGTGGCAGGCCATAGCCCGCATAATTACGAGCAATATGAACAA
CAAACCGCAGGTGAGACAGGATCAGCGTTTTAGCTGCTTCCAGATCGCCATGGTAATGCAG
CTTTTCAGCCAGCGCCCGCTCCTCGTCAGCCGACAACATCGGCCACG

DNA oligos for the simplified construction by oligo annealing
DEB3L prototype (60 oligos)
/5/ TTTATTCTTGTAATACTAGTCATCCTCGTGATGCTGGAAACAAGACCACGAGCAGGC-
CCG /3/

C - DEB3L
/5/ TCGA CGGGCCTGCTCGTGGTCTTGTTTCCAGCATCACGAGGATGACTAGTATTACAA-
GAATAAA AGCT /3/

DEB3R prototype (60 oligos)
/5/ GCTGGAAACAAGACCACGAGCAGGCCCGCGGGTTTATTCTTGTAATACTAGTCATC-
CTCG /3/

C - DEB3L
/5/ TCGA
CGAGGATGACTAGTATTACAAGAATAAACCCGCGGGCCTGCTCGTGGTCTTGTTTCCAG
AGCT /3/

**Detailed information on cell strains**

We purchased K-12 collection of *E. coli* from the Coli Genetic Stock Center in Yale University. The genetic background of the strains are as follows:

wild type (BW25113, #7636)
$F^-$, Δ(araD − araB)567, ΔlacZ4787(:: rnB − 3), λ − , rph − 1, Δ(rhaD − rhaB)568, hsdR514

ΔuvrD (JW3786-5, #10752)
$F^-$ , Δ(araD−araB)567, ΔlacZ4787(:: rnB−3), λ − , rph−1, ΔuvrD769 :: kan, Δ(rhaD−rhaB)568, hsdR514
ΔmutS (JW2703-2, #10126)
$F^-$, Δ(araD−araB)567, ΔlacZ4787(:: rnB−3), λ − , rph−1, ΔmutS738 :: kan, Δ(rhaD−rhaB)568, hsdR514

ΔmutL (JW4128-1, #10971)
$F^-$ , Δ(araD−araB)567, ΔlacZ4787(:: rnB−3), λ − , rph−1, ΔmutL720 :: kan, Δ(rhaD−rhaB)568, hsdR514

ΔmutH (JW2799-2, #10186)
$F^-$ , Δ(araD−araB)567, ΔlacZ4787(:: rnB−3), λ − , rph−1, ΔmutH756 :: kan, Δ(rhaD−rhaB)568, hsdR514

ΔuvrB (JW0762-2, #8819)
$F^-$ , Δ(araD−araB)567, ΔlacZ4787(:: rnB−3), λ − , rph−1, ΔuvrB751 :: kan, Δ(rhaD−rhaB)568, hsdR514

Δdam (JW3350-2, #11675)
$F^-$ , Δ(araD − araB)567, ΔlacZ4787(:: rnB − 3), λ − , rph − 1, Δdam − 722 :: kan, Δ(rhaD − rhaB)568, hsdR514

ΔrecA (BW26355, #7651)
$F^-$ , Δ(araD−araB)567, ΔlacZ4787(:: rnB−3), λ − , rph−1, ΔrecA635 :: kan, Δ(rhaD−rhaB)568, hsdR514

### Supplementary Discussion

#### Clan compositions are position dependent

Figure S9A shows the *s* histograms for the trimer library plotted vs. the mismatch position. The expected trimodal distribution was observable, and the dominance of C- and V-type clan peaks (*s*=0 and *s*~1, respectively) over the U-type clans indicates that most mismatches are very well repaired. Peculiarly, however, the peak corresponding to the V-type clans was not at s=1 and was instead broadly distributed yet still close to 1. Furthermore, the major V-type peak shifted to lower *s* values as the mismatch moved further away from the tracking barcode. We consistently detected this trend in all libraries examined, although the magnitude of position dependence varied between different consensus sequences. As we discuss below, this might be a phenomenon caused by the unidirectional replication of the plasmid[1] with the replication fork arriving at the mismatch region after mismatch detection and GATC site cleavage by MutH but before completion of MMR.

#### Strand asymmetry due to unidirectional replication

We initially expected to obtain trimodal s-histograms, with C- and V-peaks displaying a mirror symmetry if there are no marks distinguishing the two strands of the transformed mismatch-bearing constructs. However, this strand symmetry assumption might not hold in the presence of unidirectional replication originating from oriC [1] as well as the asymmetric distribution of the GATC motifs that are target sites for the nicking enzyme MutH. Here we would like to speculate how this effect might play a role in our system and hence explain why we observe the V-peaks in the *s* histograms to be substantially wider than C-peaks and that its peak position shifts in a reproducible and sequence-dependent manner.

Figure S10A depicts four possible cases that are possible considering the strandedness of the nick by MutH and its position with respect to the mismatch. In all four scenarios, the replication fork approaches the mismatch site from the right side, corresponding to a barcode-distal to barcode-proximal locomotion in our system. Due to the design of our constructs, the left side of the variable strand (**V**) constitutes the 5' end of the DNA, and therefore V-strands are always replicated using continuous leading strand synthesis, whereas the constant strand (**C**) is synthesized using discontinuous lagging strand synthesis mechanisms.

In Cases 1 and 2, the introduced nick is on the barcode-distal side of the mismatches and therefore the replication machinery encounters the nick before the mismatch while Cases 3 & 4 occur if the nick is generated on the barcode-proximal side of the mismatches. We expect the latter two scenario to be much more likely in our system because of the asymmetric distribution of the nearest GATC motif targeted by MutH (Fig. S10B).

In Cases 3, the introduced nick is on the V-strand, causing the replication of the leading strand to stall, whereas the intact C-strand can be replicated uninterruptedly via lagging strand synthesis. As such, the resulting clan will exclusively consist of C-strands contributing to the sharp C-peak at *s*=0.

In Case 4, the nick occurs on the C-strand, and hence the lagging strand synthesis is disrupted, while leading strand synthesis can traverse across the V-strand smoothly. As such, most of the reads in the resulting clan will originate from the V-strand, hence contributing to the V-peak in the resulting s-histogram. In contrast to the Case 3, though, the strand complementary to the C-strand can still be (partially) synthesized via Okazaki fragments and hence might be more stable against degradation than the ssDNA in Case 3. Potential replication products of the double strand break repair could result in a V-clan with incomplete domination of V-strands over C-strands.

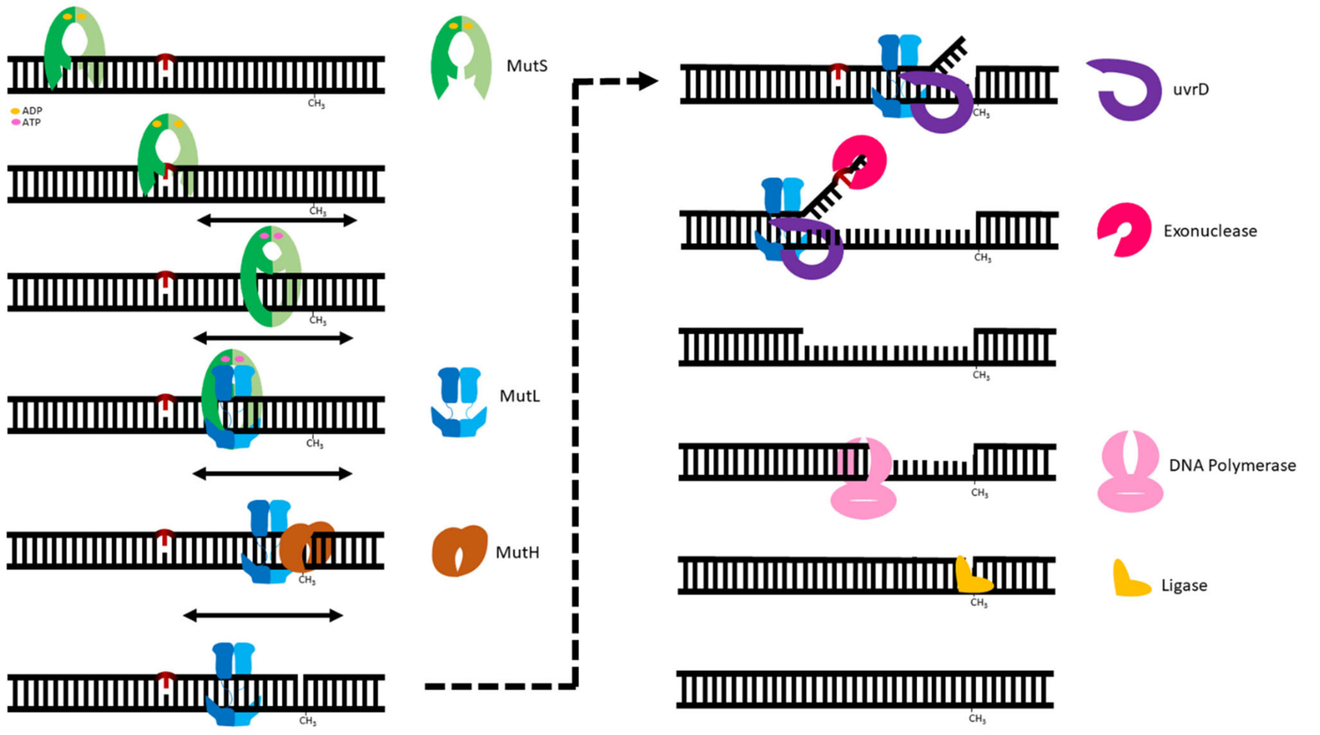


**Figure S1. Individual steps in mismatch repair in *E. coli***

*E. coli* MutS initially scans the DNA until it comes across a DNA mismatch (shown in red). MutS initially binds to the mismatch and then exchanges ADP for ATP resulting in a conformational change in MutS, resulting in a sliding clamp, where MutS will slide along the DNA to recruit and load MutL to the DNA and form a MutL sliding clamp [2]. MutL will recruit the nickase, MutH[2] which will nick the nearby hemimethylated GATC sites on the newly synthesized DNA strand. Later, MutL will recruit the helicase UvrD and the exonuclease to unwind and remove the nascent strand of DNA containing the error, respectively. DNA polymerase synthesizes DNA and DNA ligase seals the DNA to fill and repair the DNA gap and mismatch.

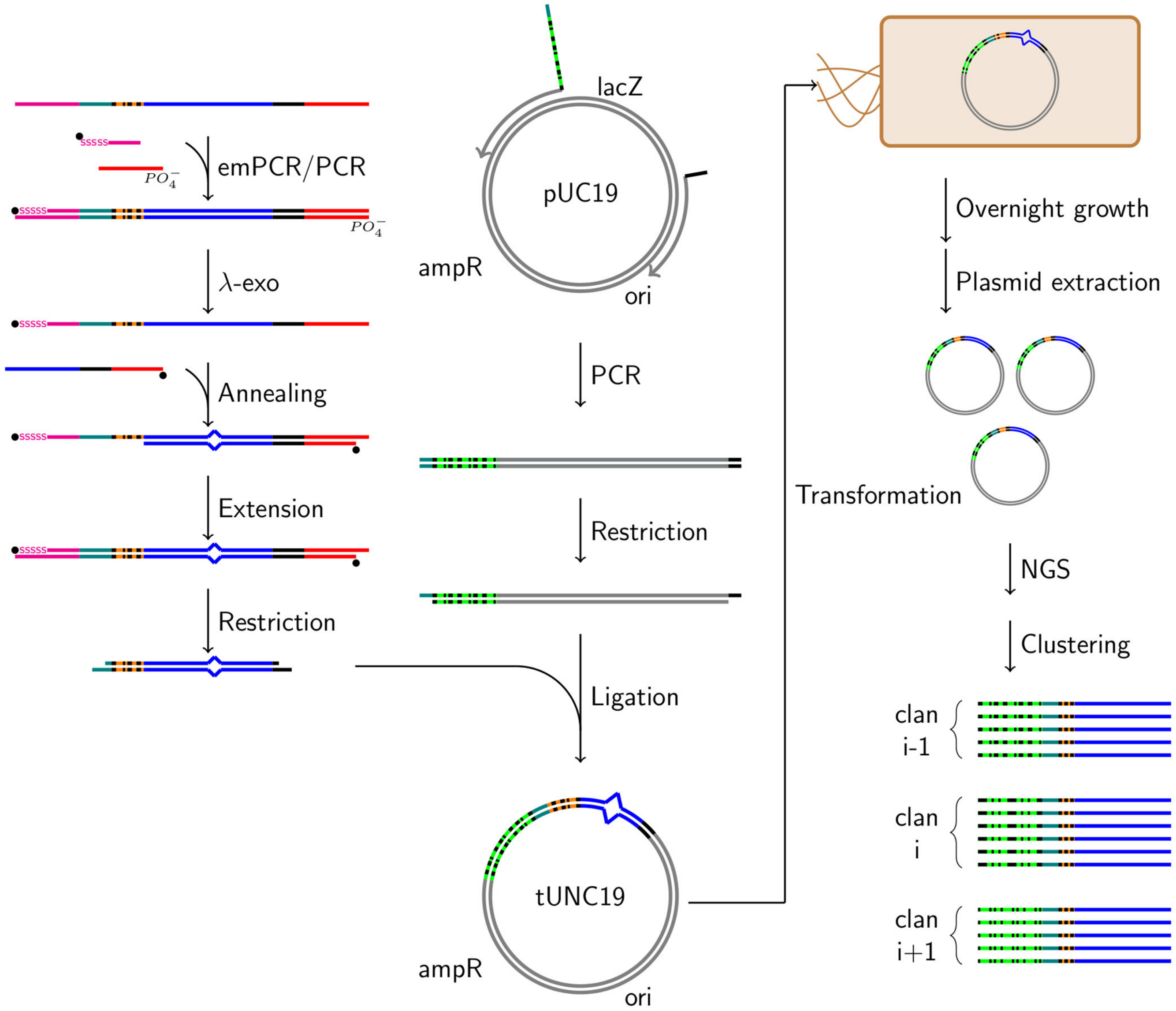


**Figure S2. Schematic summary of MMR-seq.**
An array synthesized oligo library was purchased and amplified with modified primers conferring an ability to selectively digest one of the strands. The obtained ssDNA library is annealed to an oligo of constant sequence forming a mismatch library. The mapping barcode information is copied to the other strand by primer extension. DNA barcodes were introduced to pUC19 plasmid via a PCR reaction with primers containing a random base tail that uniquely labels each individual plasmid. The resulting linear PCR product was ligated to the mismatch library after generating sticky ends with a restriction double digest reaction. The plasmid library with mismatches was transformed into *E. coli*. After multiple rounds of replication, the extracted plasmid library is sequenced. Clustering of the reads with respect to the tracking barcodes segregate DNA sequences originating from the same original plasmid, whereas the mapping barcodes are used as a lookup table for the mismatch type. The homogeneity or heterogeneity of the clans provide a means to deduce repaired and unrepaired plasmids, respectively.

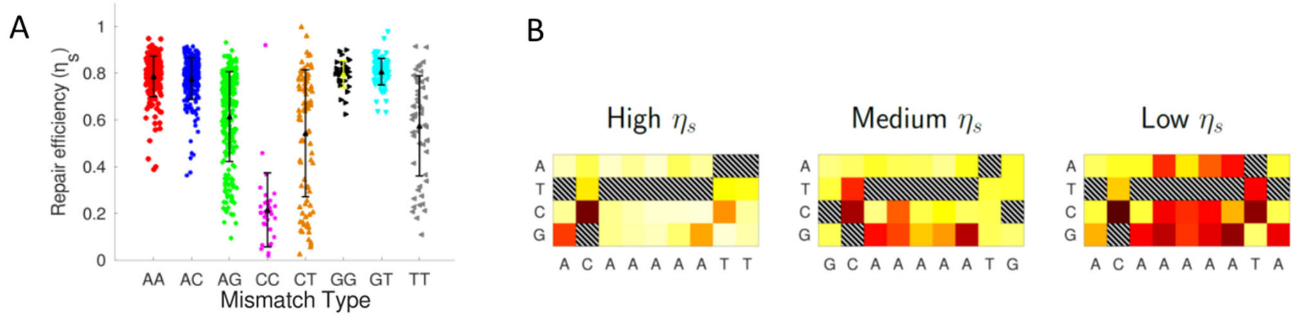


**Figure S3. MMR-seq results of partial heptamer library**

(A) Scaled repair efficiency for eight types of mismatches. Each datapoint is for a single heptamer context. Mean and standard deviation are shown in a symbol and error bar. (B) Three nonameric contexts with very different repair efficiencies even when they share the same heptameric context.

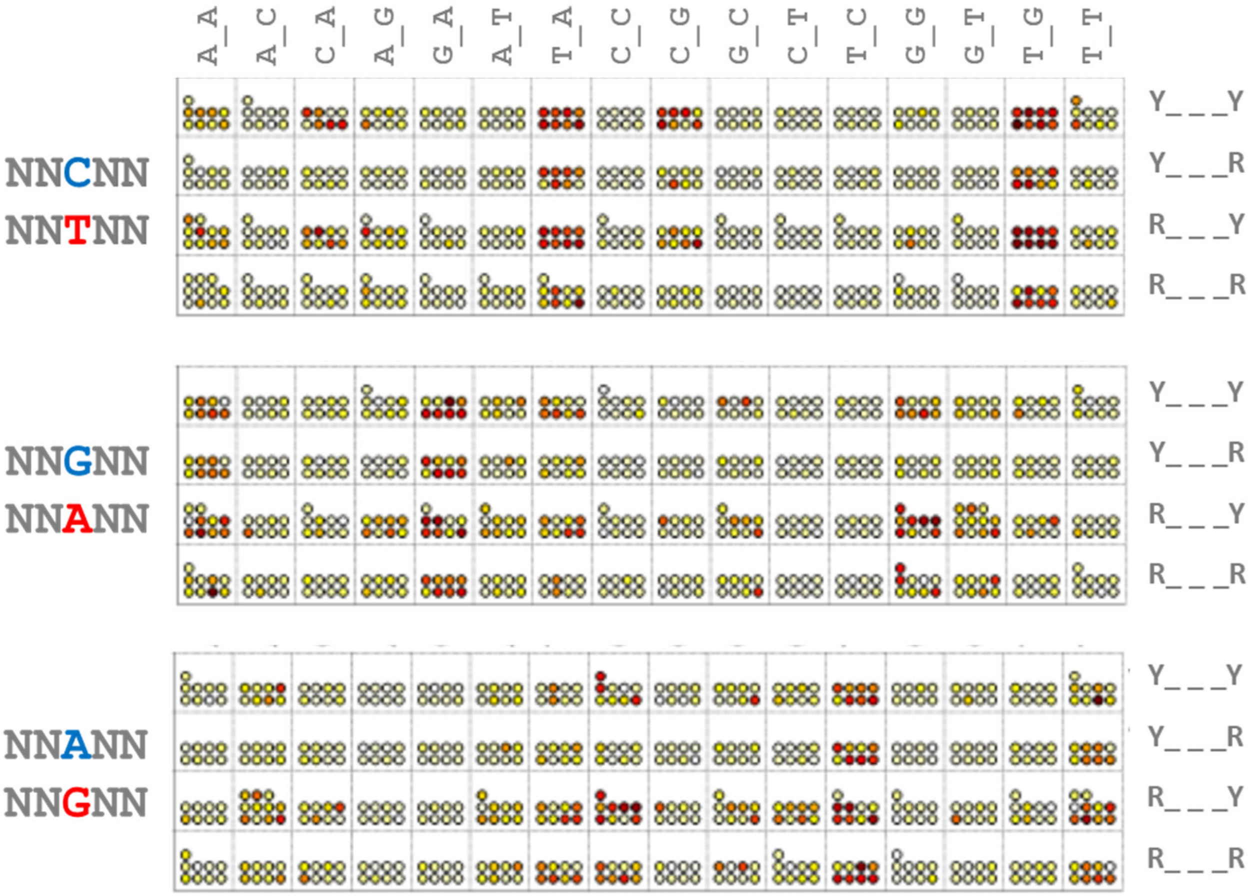


**Figure S4. Repair efficiency dependence on nearest and next nearest neighbors**
Heat map of scaled repair efficiency for CT mismatch (top), GA mismatch (middle) and AG mismatch (bottom), grouped according to the nearest surrounding nucleotides (columns) and the next nearest nucleotides (row). Y for pyrimidines (C or T) and R for purines (A or G).

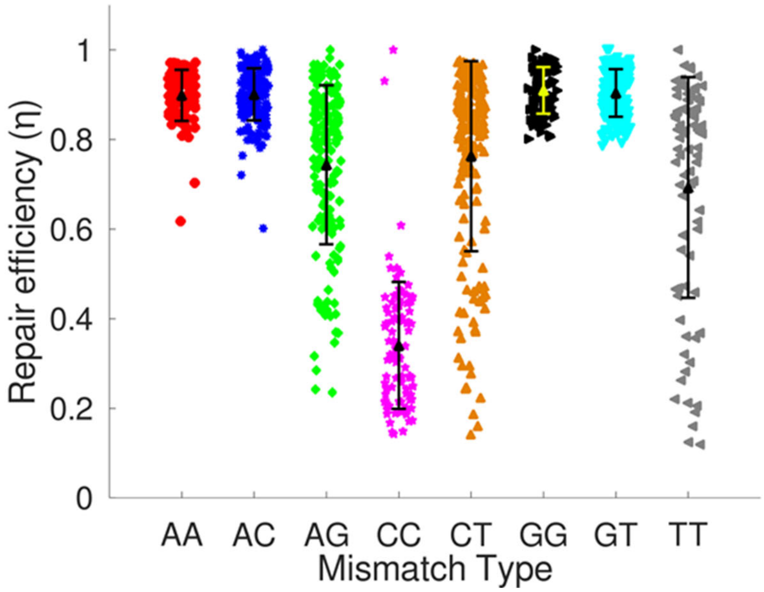


**Figure S5. MMR-seq results of *E. coli* genomic sequence library**

Repair efficiencies for eight types of mismatches obtained from three sub-libraries containing *E. coli* genomic sequences (H1, H2 and H3). Each datapoint is for a single pentamer context.

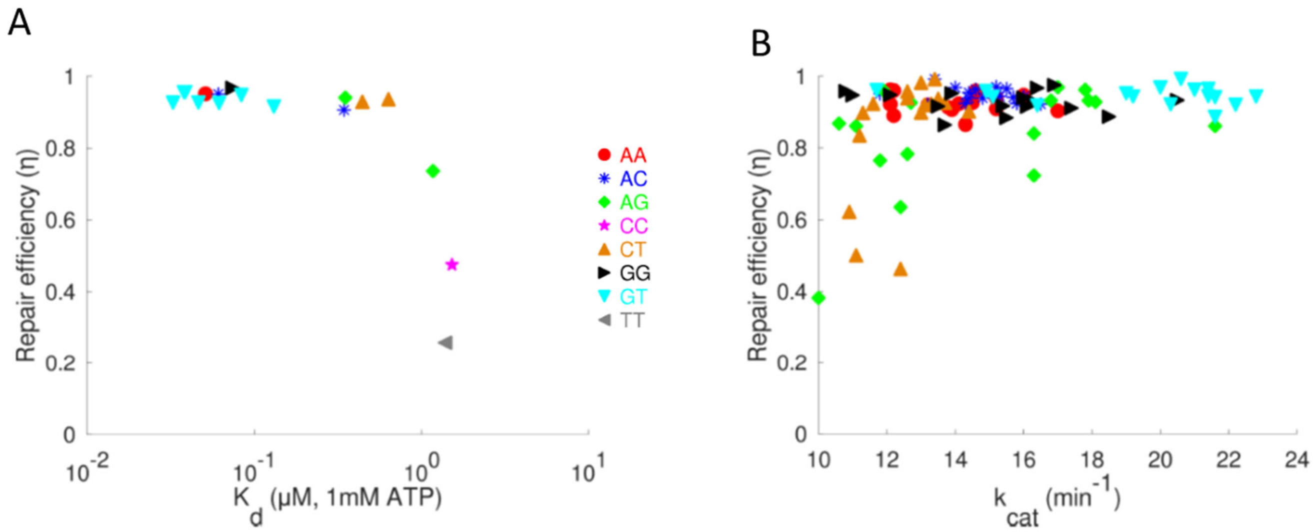


**Figure S6. Comparison to MutS biochemical data**

(A) Dissociation constant $K_d$ for MutS binding measured using surface plasmon resonance[3] vs repair efficiency in *E. coli* for the same pentamer context. Color coding for different mismatch types is shown. (B) Catalytic rate constant $k_{cat}$ for $MutS_\alpha$ ATPase activity[4] vs repair efficiency in *E. coli* for the same pentamer context.

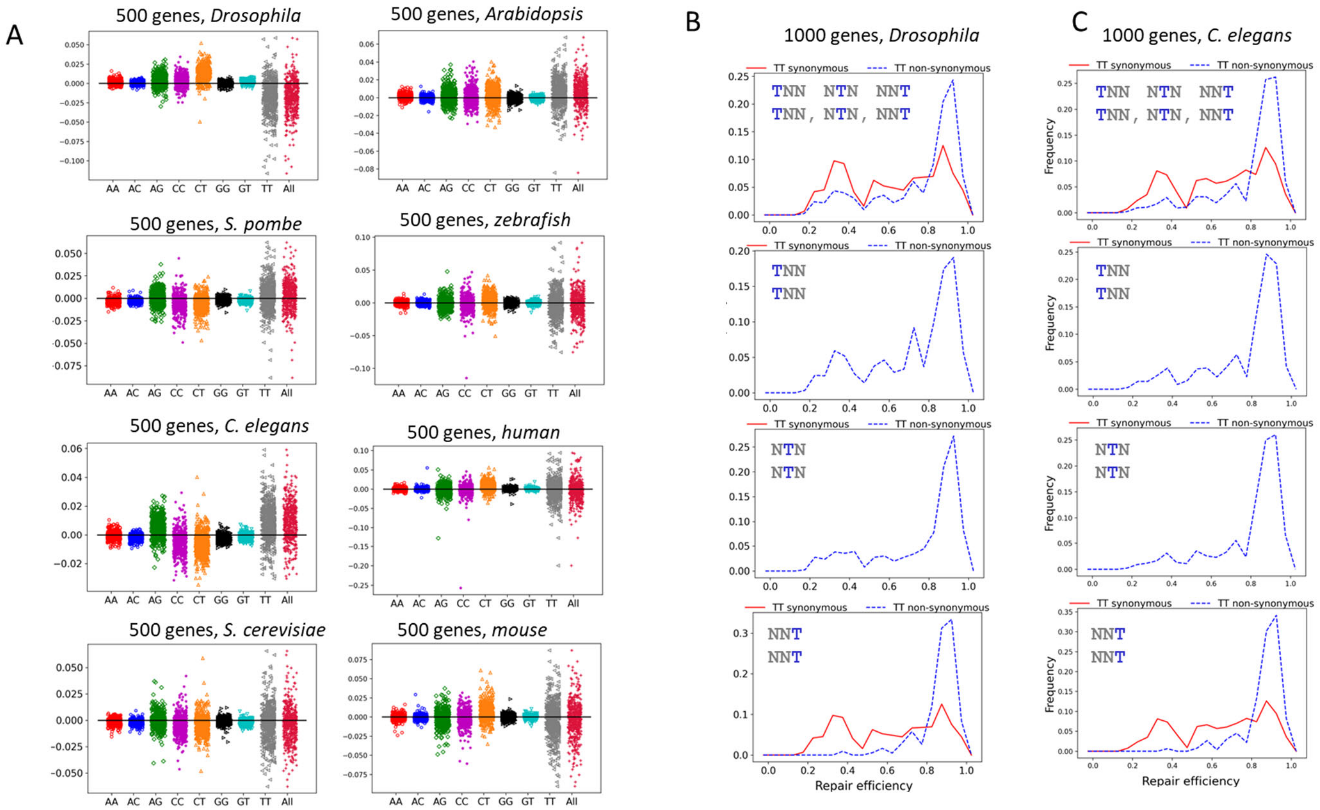


**Figure S7. Context-dependent repair and selectively enhanced mutability**
(A) Repair efficiency averaged over all possible mismatches of a given type in a protein coding sequence is predicted relative to the same gene but removing the codon bias. Contributions from TT mismatches dominate the overall behavior (in the column labeled as "All") where the exiting sequence context is more mutable compared to the case without any codon bias. Analysis was done for 500 genes each for *Drosophila*, *S. pombe*, *C. elegans, S. cerevisiae, Arabidopsis, zebrafish*, human and mouse. (B) Repair efficiency histograms for TT mismatches grouped into the cases where a failure to repair would cause a non-synonymous codon change (blue dashed) and other cases where a failure to repair would cause a synonymous codon change (red). In addition, histograms for TT mismatches in positions 1, 2 or 3 in a codon are separately plotted. 1000 genes in *Drosophila* and *C. elegans* were included.

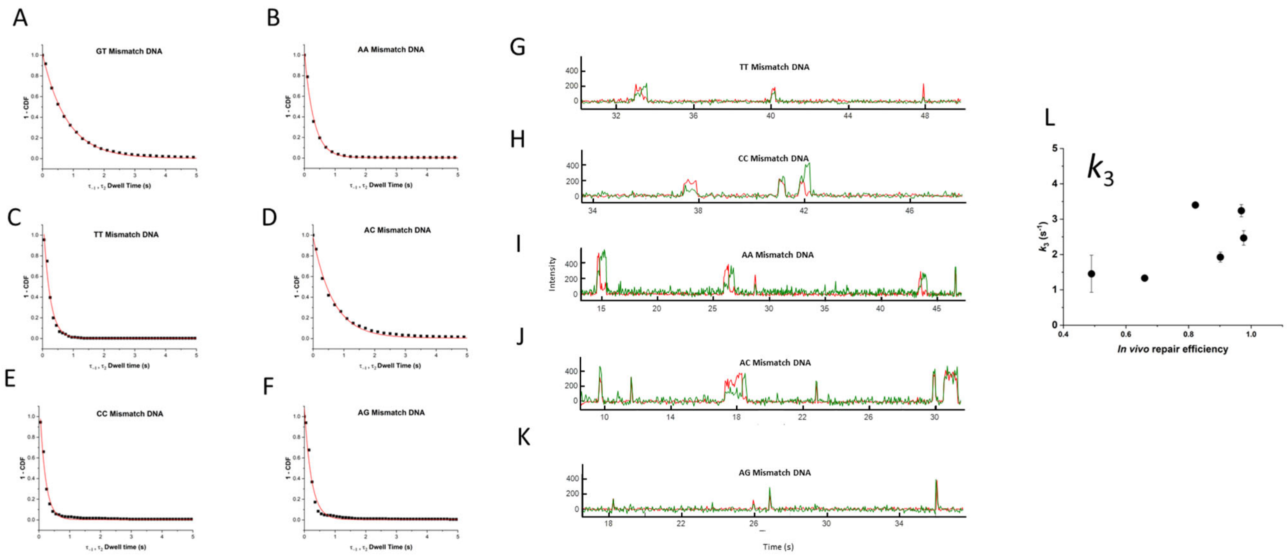


**Figure S8.** ***In vitro*** **single molecule MutS displays different binding characteristics for different mismatches**

(A-F) Cumulative distribution plots for *in vitro* MutS binding to six DNA mismatches fitted with an exponential decay curve (shown in red). (G-K) Representative single molecule traces of MutS binding to TT, CC, AA, AC and AG mismatches. (L) The rate at which MutS slides off the DNA, $k_3$, minimally contributes to the overall binding behavior of MutS to different DNA mismatches.

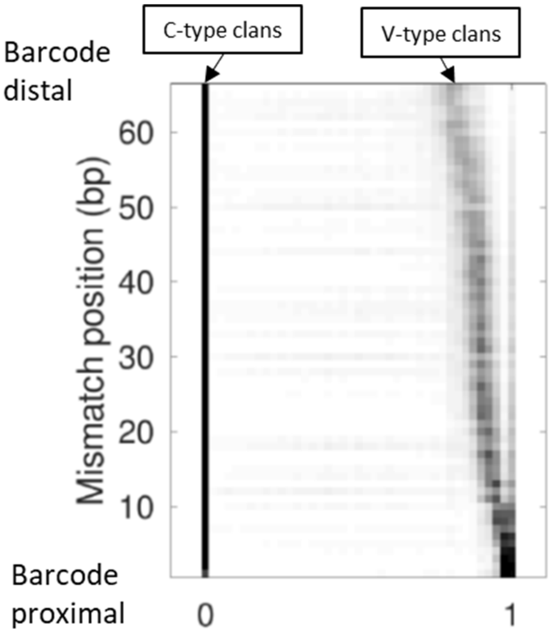


**Figure S9. Position dependence of clans repaired using the variable strand as the template**

Histograms of *s* values are displayed as a density plot (darker for higher density). A total of 198 histograms (66 positions of De Broijn trimer library times 3 types of mismatches per position) are included. The C-type clans, repaired using the constant strand as the template, stays at a single position, *s* = 0 for all cases whereas the V-type clans, repaired using the variable strand as the template, shifts to lower *s* values when the mismatch moves away from the barcodes.

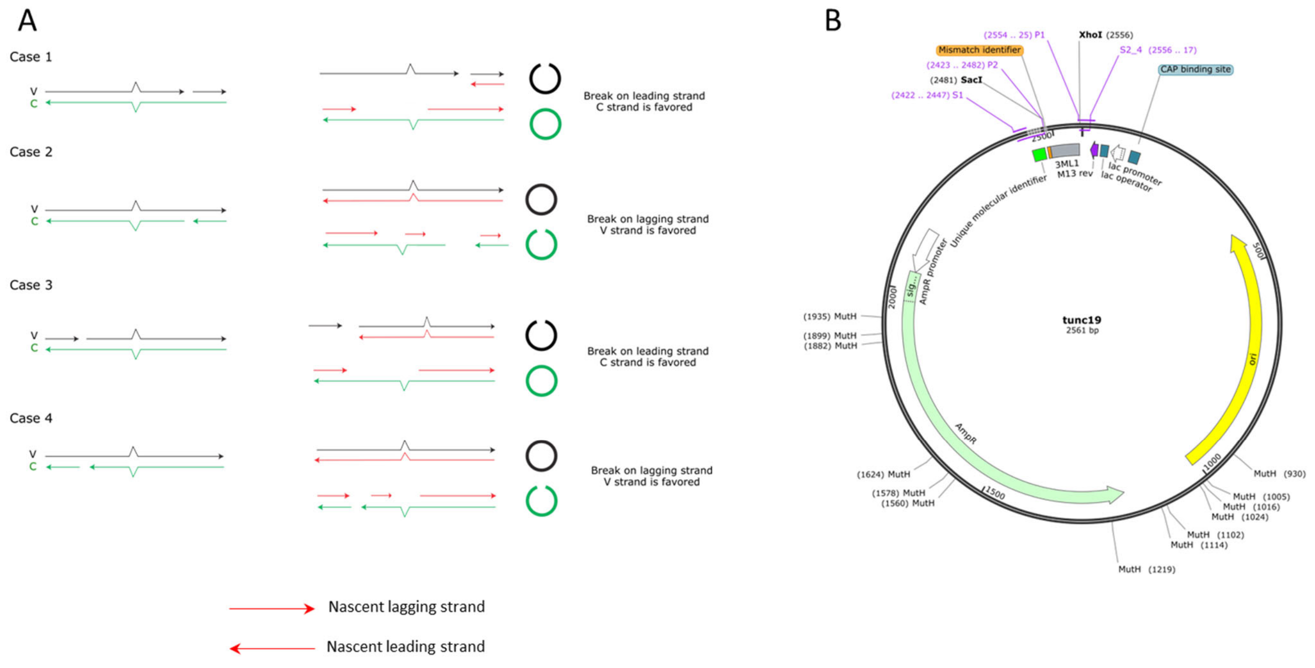


**Figure S10. Four different scenarios of mismatch repair intermediate encountered by unidirectional replication machinery**

(A) Case 1: A MutH generated nick on the leading strand template between mismatch and *OriC*. Case 2: A nick is on the lagging strand template between mismatch and *OriC*. Case 3: A nick is on the leading strand template to be encountered by the replication machinery past the mismatch as it moves to the left. Case 4. A nick is generated on the lagging strand template and will be encountered by the replication machinery past the mismatch. (B) Plasmid map shows that MutH nick sites (GATC sites) are located to favor Case 3 and Case 4.
